# The *pho1;2a’-m1.1* allele of *Phosphate1* conditions mis-regulation of the phosphorus starvation response in maize (*Zea mays* ssp. *mays* L.)

**DOI:** 10.1101/2022.03.10.483828

**Authors:** Ana Laura Alonso-Nieves, M. Nancy Salazar-Vidal, J. Vladimir Torres-Rodríguez, Ricardo A. Chávez Montes, Leonardo M. Pérez-Vázquez, Julio A. Massange-Sánchez, C. Stewart Gillmor, Ruairidh J. H. Sawers

**Affiliations:** Langebio, Unidad de Genómica Avanzada, Centro de Investigación y de Estudios Avanzados del Instituto Politécnico Nacional (CINVESTAV-IPN), Irapuato, C.P. 36824, Guanajuato, México; Department of Evolution and Ecology, University of California–Davis, One Shields Avenue, Davis, CA 95616, USA; Division of Plant Sciences, Univ. of Missouri, Columbia, MO 65211, USA; Center for Plant Science Innovation, University of Nebraska-Lincoln, Lincoln, NE, 68588, USA; Unidad de Biotecnología Vegetal, Centro de Investigación y Asistencia en Tecnología y Diseño del Estado de Jalisco A.C. (CIATEJ) Subsede Zapopan, Guadalajara, México; Department of Plant Science, The Pennsylvania State University, State College, PA, USA

## Abstract

PHO1 proteins play a central role in plant inorganic phosphorus translocation and sensing. The maize (*Zea mays* ssp. *mays*) genome encodes two co-orthologs of the *Arabidopsis PHO1* gene, designated *ZmPho1;2a* and *ZmPho1;2b*. Here, we report the characterization of the transposon-footprint allele *Zmpho1;2a’-m1*.*1*, which we refer to hereafter as *pho1;2a*. The *pho1;2a* allele is a stable derivative formed by excision of an *Activator* element from the *ZmPho1;2a* gene. The *pho1;2a* allele contains an 8 bp insertion at the point of excision that disrupts the reading frame and is predicted to generate a premature translational stop. We show that the *pho1;2a* allele is linked to a dosage-dependent reduction in transcript accumulation and a mild reduction in seedling growth that is enhanced under nutrient deficient conditions. Characterization of the shoot and root transcriptomes of seedlings segregating the *pho1;2a* mutation under different nutrient conditions revealed *pho1;2a* to have a dominant effect on patterns of transcript accumulation. Gene set enrichment analysis of the transcripts mis-regulated in *pho1;2a* mutants suggests that *Pho1;2a* functions in the fine-tuning of the transcriptional phosphate starvation response. We discuss our results with reference to possible genetic redundancy among maize *Pho1* genes and in the context of reports linking functional variation in *Pho1;2a* to agronomically important traits.

## INTRODUCTION

Phosphorus (P) is essential for crop growth and productivity. P is required for photosynthesis and respiration and is a structural component of both nucleic acids and phospholipids (Péret et al., 2011; Plaxton and Tran, 2011; Veneklaas et al., 2012; Chien et al., 2018). Elemental P is highly reactive and readily oxidized to orthophosphate (PO_4_^3-^; hereafter, inorganic phosphate, Pi) which occurs in neutral soil solution as a mixture of hydrogen phosphate (HPO_4_^2-^) and dihydrogen phosphate (H_2_PO_4_^-^). To access soil P, plants must acquire it in the form of free Pi (Schachtman et al., 1998). However, Pi has a strong tendency to react with other soil components (at low pH, predominantly iron and aluminum, at high pH, calcium and magnesium), reducing both its mobility and its availability to plants (Hinsinger, 2001; Vance et al., 2003), which ranges the concentration of plant-available P in the soil solution between 2 - 10 μM (Raghothama, 1999).

It is estimated that P availability is below the level required to realize yield in ∼70% of agricultural soils (López-Arredondo et al., 2014). For optimal growth, plants must maintain an intracellular Pi concentration of 5 - 20 mM, far higher than the concentration available in the soil (Vance et al., 2003; Raghothama, 1999). To acquire Pi against this concentration gradient, plants actively transport Pi through high-affinity transporters of the PHOSPHATE TRANSPORTER1 (PHT1) family (Rausch and Bucher, 2002; Misson et al., 2004). Once acquired, Pi is distributed around the plant through further action of the PHT1 proteins (*e*.*g*. Ai et al., 2009; Chang et al., 2019). Pi translocation is supported by the action of PHOSPHATE1 (PHO1) proteins. In *Arabidopsis*, PHO1 is indispensable for the translocation of Pi from roots to shoots, acting mainly as a Pi efflux transporter (Poirier et al., 1991; Secco et al., 2010; Bulak Arpat et al., 2012; Zhao et al., 2019; Ma et al., 2021). *PHO1* is expressed in the pericycle and root xylem parenchyma cells where it plays a role in the efflux to Pi to the xylem for translocation (Bulak Arpat et al., 2012). PHO1 proteins contain a tripartite N-terminal SPX domain and a hydrophobic C-terminal EXS domain (Stefanovic et al., 2011; Secco et al., 2012; Ried et al., 2021). The broader family of SPX-containing proteins plays an important role in Pi sensing and signaling through physical interaction with other proteins and inositol pyrophosphate molecules (Ried et al., 2021). The EXS domain is required for the subcellular localization of PHO1 to the Golgi and *trans*-Golgi networks and is necessary for Pi export (Wang et al., 2004; Stefanovic et al., 2007: Secco et al. 2012 ; Wege et al., 2016).

Plants have evolved developmental, molecular, physiological, and metabolic strategies to optimize both P acquisition and internal P use efficiency. These strategies are modulated by plant P status and collectively referred to as the P starvation response (PSR). At the molecular level, P starvation promotes widespread change at both the transcriptional and post-transcriptional levels (Rausch and Bucher, 2002; Shin et al., 2004; Chang et al., 2019; Torres-Rodríguez et al., 2021). Physiological experiments have demonstrated that plants respond both locally to the Pi concentration of the environment and systemically to their internal Pi status (Thibaud et al., 2010; Lin W-Y et al., 2014). The phosphate starvation response is tightly regulated by a transcriptional network that converges on the MYB transcription factor PHOSPHATE STARVATION RESPONSE 1 (PHR1), a master regulator of P sensing and signaling (Puga et al., 2014; Wang et al., 2014; Barragán-Rosillo et al., 2021; Ried et al., 2021; Paz-Ares et al., 2022; Bustos et al., 2010). In *Arabidopsis*, partial loss-of-function of *PHO1* (*e*.*g*. in under-expression lines or following partial complementation of *pho1* mutants by heterologous expression) uncouples low internal Pi status from the transcriptional and physiological symptoms of P starvation (Rouached et al., 2011). Wild type *Arabidopsis* plants will maintain vacuolar Pi reserves even when suffering from P starvation. Under similar conditions, plants with reduced PHO1 function fully mobilize vacuolar Pi reserves allowing for continued growth (Rouached et al., 2011). As such, the PHO1 function can be considered to link plant Pi status, the PSR, and the development of P deficiency symptoms.

*OsPHO1;2*, the functional rice ortholog of *AtPHO1*, is abundantly expressed in roots and is required for Pi translocation (Secco et al., 2010). Interestingly, *OsPHO1;2* sense transcript is associated with a *cis*-NAT. The *cis*-NAT_*OsPHO1;2*_ is upregulated under P starvation to promote the translation of the sense *OsPHO1;2* transcripts and protein accumulation through dsRNA association unfolding the sense transcript and allowing ribosome association (Jabnoune et al., 2013; Reis et al., 2021).

In maize, there are two co-orthologs to *Arabidopsis PHO1* designated *Pho1;2a* and *Pho1;2b* (Salazar-Vidal et al., 2016). The maize *PHO1* family also contains the two more distantly related paralogs *Pho1;1* and *Pho1;3*. The coding sequences of *Pho1;2a* and *Pho1;2b* are highly similar, reflecting their presumed origin during an ancient whole-genome duplication (Salazar-Vidal et al., 2016). *Pho1;2a* and *Pho1;2b* transcripts accumulate preferentially in the roots (Salazar-Vidal et al., 2016), showing similar patterns of accumulation with respect to tissue specificity and P availability, although their absolute expression level differs, with *Pho1;2b* transcript accumulation ∼30 fold greater than that of *Pho1;2a* expression (Woodhouse et al., 2021). *Pho1;2a* is associated with a putative noncoding *cis*-NAT RNA (Salazar-Vidal et al., 2016). To date, there is no evidence of *cis*-NAT transcripts associated with the maize *Pho1;2b* gene. CRISPR-Cas9 editing has suggested maize *Pho1;2a* and *Pho1;2b* to play non-redundant roles in grain filling, with edited lines showing reduced starch synthesis and small and shrunken kernels (Ma et al., 2021). However, to date, there are no reports concerning the role of maize *PHO1* genes in root-to-shoot Pi translocation or the PSR.

To study PHO1 function in maize, we previously identified insertion of an *Activator* (*Ac*) transposon into the sixth exon of the maize *Pho1;2a* gene (*pho1;2a-m1::Ac*; (Salazar-Vidal et al., 2016). The original *pho1;2a-m1::Ac* allele showed a high level of somatic instability, prompting us to generate the stable *footprint* derivative (Bai et al., 2007) *pho1;2a’-m1*.*1 (*Salazar-Vidal et al., 2016). Here, we report the characterization of *pho1;2a’-m1*.*1* mutants. We found the *pho1;2a’-m1*.*1* mutation to have only subtle effects on plant growth and no statistically significant impact on total P concentration in the leaf. We characterized root and leaf transcriptomes in a stock segregating the *pho1;2a’-m1*.*1* mutation under different nutrient regimes, and observed up-regulation of several hundred transcripts on both heterozygous and homozygous mutant plants. Many of the transcripts upregulated in plants carrying *pho1;2a’-m1*.*1* were also induced by low P availability in wild-type plants, suggesting a role for *Pho1;2a* in fine-tuning of the transcriptional P starvation response.

## MATERIALS AND METHODS

### Plant material and genotyping

Analysis of the wild-type siblings of the mutant segregants described here was reported previously in Torres-Rodríguez et al., 2021. Three self-pollinated sibling families segregating for the *pho1;2a’-m1*.*1* mutation in the *Zea mays* ssp. *mays* var. W22 background were used for the experiments (Salazar-Vidal et al., 2016). Plants were genotyped as previously reported (Salazar-Vidal et al., 2016) with small modifications. Briefly, DNA was extracted from a punch of cotyledon tissue and a fragment was amplified spanning the position of the *pho1;2a’-m1*.*1* 8 bp insertion using the primers MS002 and MS126 (Supplemental Table 1). PCR was performed using Kapa Taq DNA polymerase (Kapa Biosystems), under the following cycling conditions: denaturation at 95 °C for 5 min; 35 cycles of 95 °C for 30 sec, 60 °C for 40 sec, 72 °C for 30 sec; final extension at 72 °C for 5 min. PCR products were digested with BseYI enzyme (New England BioLabs) according to the manufacturer’s protocol and analyzed by gel electrophoresis to determine the genotype.

### Growth conditions

Plants were grown under a greenhouse system as previously described (Torres-Rodríguez et al., 2021). Sand was used as substrate with nutrient conditions maintained by use of a combination of solid-phase P buffer (loaded with 209 μM KH_2_PO_4_ for high P treatments and 11 μM KH_2_PO_4_ for low P treatments; Lynch et al., 1990) and fertilization with modified Hoagland solution (5 mM KNO_3_, 0.25 mM Ca (NO_3_)_2_, 2 mM MgSO_4_, 1 mM KH_2_PO_4_, 20 μM FeC_6_H_6_O_7_, 9 μM MnSO_4_, 1.2 μM ZnSO_4_, 0.5 μM CuSO_4_, 10 μM Na_2_B_4_O_7_, 0.008 μM (NH_4_)_6_Mo_7_O_24_ (Hoagland and Broyer, 1936)). For low N treatments, Hoagland was adjusted by substitution of KNO_3_ and Ca(NO_3_)_2_ with KCl and CaCl_2_, respectively (Reddy et al., 1996; Escobar et al., 2006; Baxter et al., 2008). For low P treatments, Hoagland P concentration was adjusted by substitution of KH_2_PO_4_ with KCl (Diepenbrock, 1991). Hoagland solution was applied at 1/3 strength with final N/P concentrations in the different treatments as follows: Full treatment - 1750 μM NO_3_□, 333μM PO□^3^□; LowN - 157.5 μM NO_3_□, 333μM PO□^3^□; LowP - 1750 μM NO_3_□,10 μM PO□^3^□; LowNP - 157.5 μM NO_3_□, 10μM PO□^3^□.

For growth evaluation plants were grown in PVC tubes (15 cm diameter; 1 m tall) until 40 days after emergence (DAE). Tubes were filled with ∼17 liters of washed sand. In the upper third of the tube, sand was amended with 1.5 % of the full or low P loaded solid-phase P buffer. Four seeds were selected at random from segregating seed stock, imbibed and planted at 4 cm depth per tube. The experiment was established over four planting dates spaced at four-day intervals. On each day, nine tubes were established per nutrient treatment, arranged as a 3×3 group. The resulting 16 groups were arranged in a latin square with respect to nutrient treatment, although row number was confounded with planting date. Plants were thinned to a single plant a week after emergence and the retained plants were genotyped for the *pho1;2a’-m1*.*1* mutation. Plants were irrigated with distilled water up until 10 DAE after which Hoagland treatments were applied, at a rate of 200 ml every third day. Plants were evaluated every five days by non-destructive measurement of *stem width, leaf number, leaf length*, and *leaf width* for each fully expanded leaf. *Leaf area* was calculated as green leaf length x leaf width x 0.75 (Francis et al., 1969). *Total leaf area* was the sum of all leaves per plant. At 40 DAE, plants were carefully removed from the tubes, minimizing damage to the root system, washed in distilled water, and dried with paper towels. The cleaned root system was placed in a water-filled tub and photographed using a digital Nikon camera D3000. Raw images were individually processed using Adobe Photoshop CC (Version 14.0) to remove the background and maximize contrast between foreground and background non-root pixels. Processed images were scaled and analyzed using GiA Roots software (Galkovskyi et al., 2012) as previously described (Torres-Rodríguez et al., 2021). After photography, the root system was divided into segments corresponding to increments of 15 cm depth (numbered 1 to 6) and segments weighed individually. Shoot and root tissue were dried for one week at 42 °C using a drying oven before taking dry weights and collecting tissue for element quantification. Trait description is provided in Supplemental Data Set 1A.

For transcriptome analysis, plants were grown in shorter PVC tubes (15 cm diameter; 50 cm tall). Substrate and nutrient treatments were the same as for the large tube experiment with the top 30 cm of the tube amended with 1.5 % solid-phase P buffer. At 25 DAE, the whole plant was harvested, separating the total root system, stem and leaves. Leaf and root tissue for gene expression were immediately frozen in liquid nitrogen and stored at -80 °C. Samples were homogenized with cooled mortar and pestle and aliquoted under liquid nitrogen for RNA extraction and transcriptome analysis.

For quantification of *Pho1;2a* and *Pho1;2b* transcripts, a third experiment was carried out with the same growth set-up as the RNA-Seq experiment. In this experiment, two treatments were applied, Full nutrients and LowP without the use of solid-phase P buffer (alumina-P). Plants were harvested at 25 DAE as described above.

### Quantification of Pho1;2a and Pho1;2b transcripts

Total RNA was extracted using Trizol protocol (Invitrogen), and cDNA was synthesized using SuperScript II reverse transcriptase (Invitrogen) after DNase I treatment (Invitrogen) following the manufacturer’s protocol. qRT-PCR was performed using SYBR Green I-based real-time PCR reagent with the LightCycler® 480 Instrument (Roche) using the following program: 95 °C for 5 min, followed by 40 cycles of 95 °C for 15 sec; 60 °C for 20 sec; 72 °C for 20 sec. The *ZmCDK* gene (GRMZM2G149286) was used as a constitutive reference gene (Lin F et al., 2014). Relative expression was calculated as 2ΔCt, where ΔCt = (Ct of constitutive gene − Ct of target gene). Three biological replicates with three technical replicates were analyzed. Statistical was performed in R (Robinson et al., 2010; McCarthy et al., 2012; R Core Team, 2019) using R/stats::lm in the model gene expression ∼ genotype * treatment followed by Tukey’s HSD test using R/stats::TukeyHSD. Primers used are indicated in Supplemental Table 1.

### Determination of elemental concentration by inductively coupled plasma mass spectrometry

Ion concentration was determined as described previously by (Ramírez-Flores et al., 2017). Briefly, dry leaf samples were analyzed by inductively coupled plasma mass spectrometry (ICP-MS) to determine the concentration of twenty metal ions. Tissue samples were digested in 2.5 mL concentrated nitric acid (AR Select Grade, VWR) with an added internal standard (20 ppb In, BDH Aristar Plus). The concentration of the elements Al, As, B, Ca, Cd, Co, Cu, Fe, K, Mg, Mn, Mo, Na, Ni, P, Rb, S, Se, Sr, and Zn were measured using an Elan 6000 DRC-e mass spectrometer (Perkin-Elmer SCIEX) connected to a PFA microflow nebulizer (Elemental Scientific) and Apex HF desolvator (Elemental Scientific). A control solution was run for every tenth sample to correct for machine drift both during a single run and between runs.

### Statistical analysis of plant growth and ionomic data

For plants grown to 40 DAE, traits were obtained from 133 unambiguously genotyped individuals across the wild-type (wt), heterozygous (ht) and homozygous *pho1;2a-m1*.*1* (mu/mt) genotypes, four nutrient treatments (Full, LowN, LowP and LowNP), and four planting dates (Supplemental Data Set 1B). Traits included direct measurements and derived values (*e*.*g*., total leaf surface area or biomass totals). Non-destructive measurements were taken from 10 DAE and repeated at 5-day intervals during the experiment. Destructive measurements were made for all 133 individuals at harvest. The data set included element concentrations determined by ICP-MS and root architectural traits extracted by image analysis, as described above.

Statistical analysis was performed in R (R Core Team, 2022). Our focus was on differences between genotypes and we performed separate analyses for each trait/treatment combination to limit model complexity. Traits were classified as dynamic (repeated measures) or endpoint (traits collected at harvest) and analyzed separately, adjusting for multiple testing within each group. Ionomic element concentrations were also analyzed separately. For visualization, we used R/stats::lm to fit the model *trait value* ∼ 0 + *genotype* + *planting date* + *error* for each trait/treatment combination, extracting model coefficients and standard errors for plotting. For dynamic traits, we performed pairwise comparisons of wt, ht and mu curves (trait over time) for each trait/nutrient combination using R/statmod::compareGrowthCurves (Baldwin et al., 2007) under the meanT function, using 10,000 permutations for p-value estimation. p-values were adjusted using Holm’s method with R/stats::p.adjust, both within each trait/treatment combination and globally. For endpoint analysis, trait values were square root transformed to improve normality and modeled as *trait value*^*1/2*^ ∼ *planting date* + *genotype* + *error* for each trait/treatment combination, extracting the *genotype* p-value from the associated ANOVA table. Endpoint p-values were adjusted for multiple testing using Holm’s method with R/stats::p.adjust, applied separately to growth, GiA Roots and ionomic data sets. Where the treatment effect was significant (adjusted p < 0.05), we used R/agricolae::HSD.test (de Mendiburu, 2020) to apply pairwise Tukey HSD tests at *α* = 0.1 to identify differences between genotypes.

### RNA extraction and transcriptome sequencing

Transcriptome analysis was performed for each genotype (wt, ht, mt) in the four treatments (Full, LowN, LowP, and LowNP) from roots and leaves tissues represented by two biological replicates. RNA extraction and library generation was performed by Labsergen (Laboratorio de Servicios Genómicos, Langebio). Leaf libraries were sequenced using Illumina HiSeq4000 high-throughput sequencing technology at the Vincent J. Coates Genomics Sequencing Laboratory (UC Berkeley). Root libraries were sequenced by Labsergen (Laboratorio de Servicios Genómicos, Langebio) on the Illumina NextSeq 550 equipment. Transcriptome data are available in the NCBI Sequence Read Archive under study SRP287300 at https://trace.ncbi.nlm.nih.gov/Traces/sra/?study=SRP287300. A separate analysis of the libraries generated from wild-type segregants was published previously (Torres-Rodríguez et al., 2021).

### Analysis of differential gene expression

RNA sequencing reads were aligned to cDNA sequences from the AGPV3.30 version of the maize genome available at Ensembl Plants (ftp://ftp.ensemblgenomes.org/pub/plants/release-30/) using kallisto (version 0.43.1; Bray et al., 2016). Transcript abundance data were pre-processed using R/tximport (Soneson et al., 2015) with gene-level summarization. Transcript counts were analyzed with a GLM approach using R/edgeR (Robinson et al., 2010; McCarthy et al., 2012; R Core Team, 2019) and R/limma (Ritchie et al., 2015). To explore the impact of the *pho1;2a’-m1*.*1* mutation on gene regulation, we compared genotypes in separate analyses for each tissue (root or leaf) and treatment (Full, LowN, LowP, LowNP) combination (condition). We then pooled information across all analyses using R/mashr (Urbut et al., 2019) to generate the final list of differentially expressed genes (DEGs). Initial analyses were performed by generating a series of DGEList objects, one for each condition, each list containing data for wild-type, heterozygous and *pho1;2a’-m1*.*1* plants. Normalization matrices and offsets were calculated for each DGEList as described in the R/edgeR documentation. Genes with fewer than 10 counts per million (CPM) in all, or all but one, of the libraries in a given DGEList were removed before normalization with R/edgeR::calcNormFactors. Data were prepared for linear modeling using R/limma::voom with no additional normalization. Global patterns of gene expression within each condition were visualized with R/edgeR::plotMDS. We estimated the effect of the genotype (GEN) at *Pho1;2a* for each condition using the linear model:

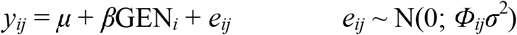

where *y*_*ij*_ is the normalized log_2_CPM value for a given gene in a given library, *μ* is the intercept for that gene in a particular condition, *β* is the effect of the genotype, and *e*_*ij*_ is the model residual, which is assumed to be independent for each *y*_*ij*_ and to have a variance proportional to *Φ*_*ij*_, the empirical weight factor calculated by voom. We fitted models using R/limma::lmfit, sent the output to R/limma::eBayes, and extracted the estimates of *β* for both heterozygous and homozygous effects, along with their corresponding standard errors, calculated as the product of the standard deviation and the residual for each gene. We generated separate objects for heterozygous and homozygous effects, each containing effect estimates and standard errors for each gene in each of the eight conditions. Where we had no estimate for a given gene in a given condition (*i*.*e*. because of filtering for low expression), we set the estimate to 0 and the standard error to 1000. Estimates and standard errors were passed to R/mashr::mash using a canonical covariance matrix. For each gene, we extracted posterior estimates for heterozygous and homozygous mutant effects in each condition, along with standard deviations and local false sign rates (lfsr). For any given gene, we reset the posterior estimate to 0 for any condition for which the value had originally been set to NA because of low expression. We combined heterozygous and homozygous mutant effects into a single object and selected genes with posterior effects > |1| and lfsr < 0.01 as differentially expressed genes (DEGs). Genes called as DEG in at least one condition for either heterozygous or homozygous effect were retained in our set of 1, 100 *Pho1;2a*-regulated genes (Supplemental Data Set 2A). Gene annotation was assigned as previously described (Gonzalez-Segovia et al., 2019). We used posterior effects to visualize differential expression between genotypes. To visualize expression differences between conditions, we ran a second linear model using all samples, considering each combination of condition and genotype as a different level of a single grouping variable and extracting coefficients (Supplemental Data Set 2A). To investigate P signaling, the response of the 1, 100 *Pho1;2a*-regulated genes were considered with respect to LowP in the roots and leaf of wild-type and mutant plants (Supplemental Data Set 2B, C). Gene ontology (GO) analysis was performed with BiNGO (v. 3.0.3; (Maere et al., 2005) in Cytoscape (v. 3.7.2) using the set of 1,100 *Pho1;2a*-regulated genes in the context of 39468 maize B73 RefGen_v3 gene models. Significant GO terms were obtained using the hypergeometric test with the Benjamini & Hochberg FDR correction and a significance level of 0.05 (Supplemental Data Set 2D).

## RESULTS

### The *pho1;2a’-m1.1* mutation introduces a premature stop codon

The maize *pho1;2a’-m1*.*1* allele (hereafter, *pho1;2a*) contains an 8 bp (CTGCCCAG) insertion at the point of *Ac* excision from the progenitor allele *pho1;2a-m1::Ac* (Fig. 1A; (Salazar-Vidal et al., 2016). The 8 bp insertion shifts the reading frame relative to the wild-type gene and generates a premature stop codon after residue 419 (based on the W22 gene model Zm00004b022763_T001), resulting in a mutant protein that is predicted to be truncated between the SPX and EXS domains (Fig. 1A). In *Arabidopsis*, the EXS domain is crucial for subcellular localization of PHO1, Pi export and root-shoot signaling (Wege et al., 2016). We predict that the *pho1;2a* mutation knocks-out native protein function, although we cannot exclude the possibility that a novel truncated SPX domain-containing product is produced (Fig. 1A). Our *pho1;2a* allele is similar to a recently reported CRISPR-Cas9 editing event that deleted 2 bp in the sixth exon of *Pho1;2a*, shifting the reading frame after residue 391 and generating a premature stop codon at residue 627 (based on the W22 gene model Zm00004b022763_T001; Ma et al., 2021).

**Figure 1.**
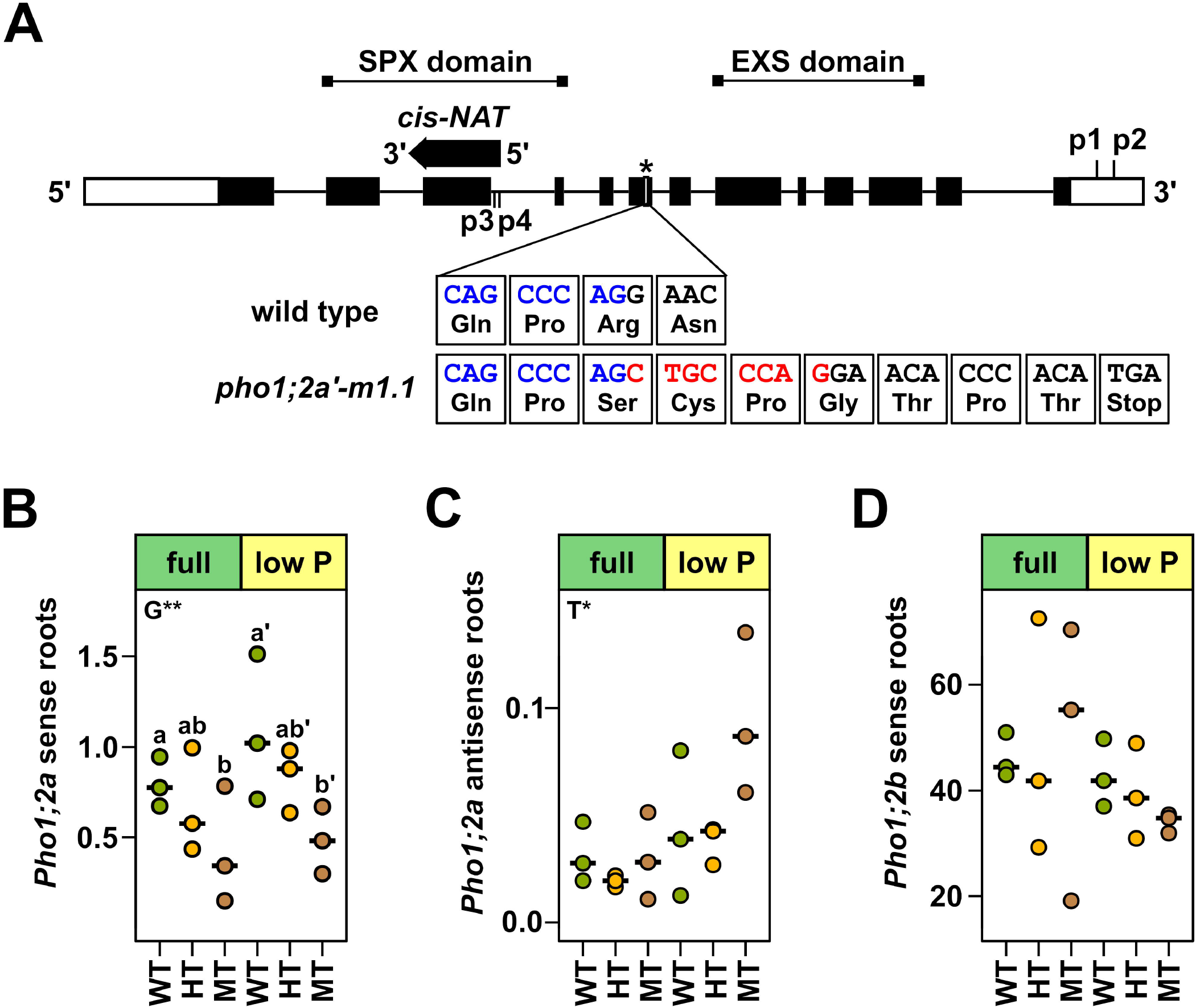
The *pho1;2a’m1.1* mutation introduces a shift in reading frame and a premature stop codon. A) Structure of the *ZmPho1;2a* gene showing the site of transposon insertion (*) in the allele *pho1;2a-m1::Ac*. The DNA sequence surrounding the insertion site in the wild type and the *pho1;2a’-m1*.*1* footprint allele is shown below, along with predicted protein sequence. Sequence adjacent to the insertion site is shown in blue. Eight base pairs of additional sequence generated at *Ac* insertion and retained following *Ac* excision are shown in red. The DNA and predicted protein sequences are based on the v2.0 of the W22 genome (Zm00004b022763). Black rectangles represent exons, black lines represent introns, white rectangles represent UTRs. The black arrow indicates the position of a *cis*-Natural Antisense Transcript. The approximate positions of regions encoding the conserved SPX and EXS protein domains are marked. P1, P2 indicate the positions of primers used to quantify *Pho1;2a* sense transcripts in RT-qPCR analysis. P3, P4 indicate primers used to quantify *Pho1;2a* antisense transcripts. B) Quantification of *Pho1;2a* transcripts in the roots of wild type (WT, green), heterozygous (HT, yellow), and homozygous mutant (MT, brown) individuals of a family segregating the *pho1;2a’m1*.*1* mutation, grown under sufficient (Full) and or low phosphate (low P) treatment. RT-qPCR was used to quantify relative expression in three individuals per condition. Transcript accumulation is shown relative to the maize CDK gene (GRMZM2G149286). C) and D), as B, showing quantification of *Pho1;2a cis-NAT* and *Pho1;2b* transcripts, respectively. In B-D, significant differences among genotypes (G) or treatments (T) are indicated as *, p < 0.05; **, p < 0.01; ***, p < 0.01.

To evaluate the effect of the *pho1;2a* mutation on the *Pho1;2a* gene, we quantified *Pho1;2a* transcripts in a *pho1;2a* segregating stock with quantitative PCR. We included a primer set designed to amplify a previously described putative anti-sense transcripts (Salazar-Vidal et al., 2016). We grew plants under both full-nutrient and low P conditions (see Materials and Methods) and assessed transcript levels at 25 days-after-emergence (DAE; Fig. 1B-D). Although we could detect *Pho1;2a* sense transcripts in all genotypes under both P conditions, there was a dosage-dependent reduction in accumulation associated with the *pho1;2a* allele (Fig. 1B). Our quantitative PCR primers were located in the 3’-UTR, indicating that full length transcripts are produced from *pho1;2a*. We did not detect a statistically significant effect of *pho1;2a* on the putative *Pho1;2a* antisense transcript, although there was some evidence of increased accumulation in homozygous *pho1;2a* mutant plants under low P (Fig. 1C). Consistent with a previous report, the level of accumulation of *Pho1;2b* sense transcripts in the roots was greater than that of *Pho1;2a* in wild-type plants (Fig. 1D). We did not detect any effect of *pho1;2a* on *Pho1;2b* transcript accumulation.

### The *pho1;2a* mutation is linked a reduction in seedling growth that is intensified under low nitrogen availability

There was no obvious phenotype associated with *pho1;2a* during generation and propagation of homozygous stocks in the field (*c*.*f. pho1* mutants in *Arabidopsis*, rice or tomato (Poirier et al., 1991; Secco et al., 2010; Zhao et al., 2019). Despite the molecular similarity of our *pho1;2a* allele to previously reported CRISPR-Cas9 events, we did not see a linked clear shrunken kernel phenotype (Ma et al., 2021). We individually weighed all the kernels from *pho1;2a* segregating stocks that were planted for seedling characterization. Based on the seedling genotype data, there was no effect of the embryo/endosperm genotype on kernel weight (*n* = 133; means: wt = 197 mg, ht = 194 g, mt = 189 mg; Kruskal-Wallis p = 0.15). We also compared 100 kernel-weight of a homozygous *pho1;2a* stock to that of a wild-type sibling family. Here, genotype was significant, although potentially acting as a maternal effect (*n* = 5; means: wt = 22.5g, mu = 20.2 g; Kruskal-Wallis p = 0.01).

The SPX domain protein family is important in P nutrition and in crosstalk between P and Nitrogen (N) signaling pathways (Hu et al., 2019; Ueda et al., 2020; Torres-Rodríguez et al., 2021). We speculated that the *pho1;2a* phenotype would be enhanced under nutrient deficiency. We used controlled greenhouse conditions to characterize seedlings from a stock segregating the *pho1;2a* mutation under different levels of N and P availability (Full, LowN, LowP and combined LowNP. See Materials and Methods). We followed plant growth by manual measurement of green leaf area (LA) every 5 days, starting at 10 DAE to 40 DAE, followed by endpoint measurements at harvest at 41 DAE. The first two leaves were fully expanded when measurements began at 10 DAE.

We observed a subtle reduction in seedling growth in *pho1;2a* mutants (Fig. 2, Supplemental Figure 1; Supplemental Data Set 1). Under Full and LowN conditions, LA of leaf (L)1 was smaller in homozygous mutants and heterozygotes than in wild-type siblings at 10 DAE, with evidence of more rapid senescence in homozygous mutants and heterozygotes, especially in Full conditions (Fig. 2A, Supplemental Figure 1; Supplemental Data Set 1C). We did not observe differences in L1 among genotypes under LowP or LowNP. L3 was initiated during the experiment, with plants reaching L8 or L9 by 40 DAE. Leaf growth was reduced in homozygous mutants and heterozygotes compared with wild-type from L4 onwards under LowN, and from L6 under LowP (Fig. 2A, Supplemental Figure 1; Supplemental Data Set 1C). Statistical support for growth effects was limited, with only effects on L1 and L2 under Full and L1-5 under LowN being significant at 5%. Significant differences in Total LA (TLA) were observed under LowN and robust to adjustment for multiple testing across traits and treatments (Fig. 2B). In all significant cases, homozygous mutants and heterozygotes were equivalent and different to wild-type, with the exception of L1 and L2 under LowN where heterozygotes followed wild-type growth. Overall, seedling analysis indicated a dominant effect of the *pho1;2a* mutation on vegetative growth that was enhanced under LowN availability. LA measurements were mirrored by endpoint shoot biomass (Supplemental Figure 2 - SFW LowN; Supplemental Data Set 1D).

**Figure 2.**
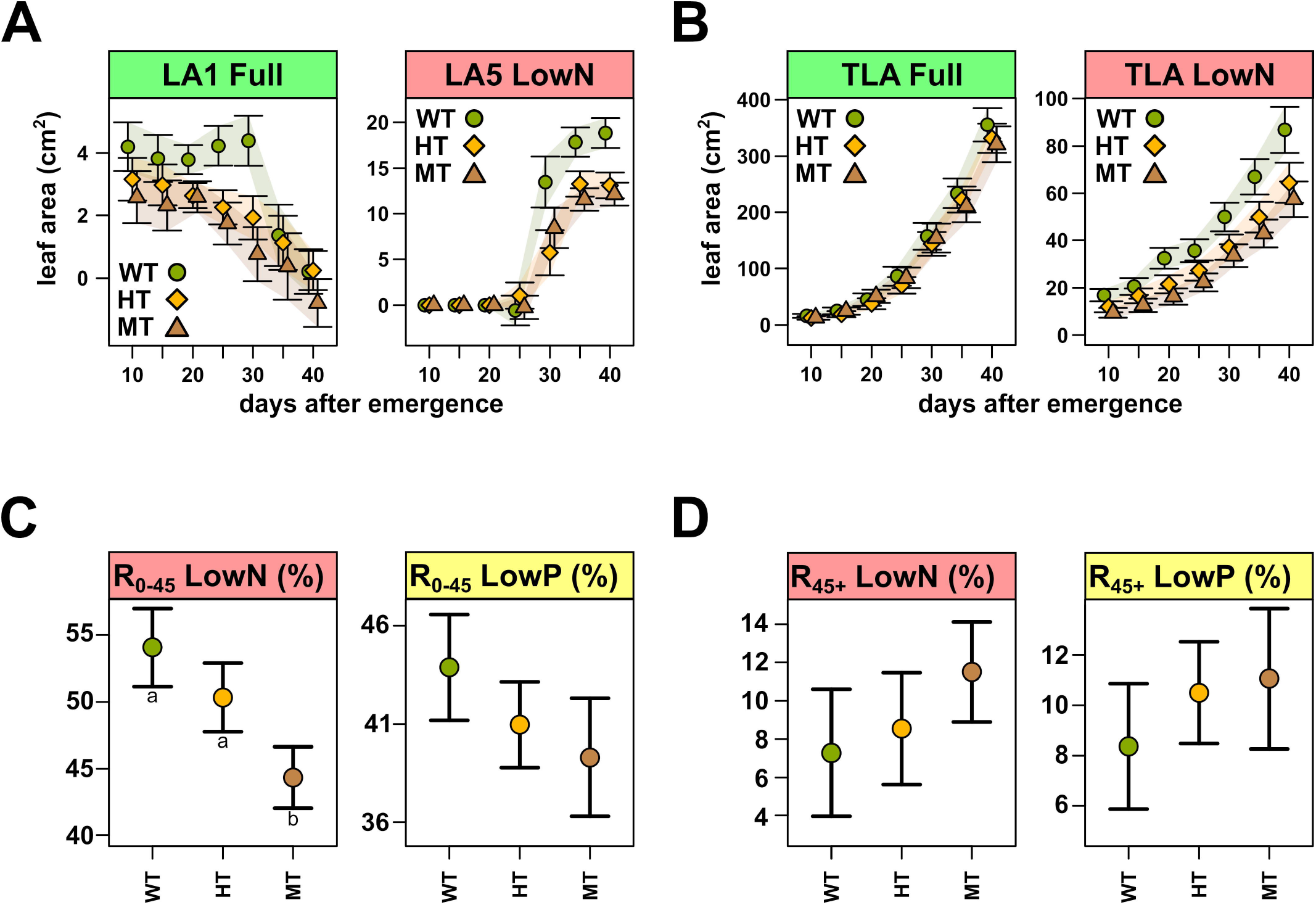
Seedling growth is reduced in *pho1;2a’m1.1* mutants. A) Fully expanded green area (cm^2^) of first (LA1) and fifth (LA5) leaves from 10 days-after-emergence (DAE) until day 40. The number of fully expanded green leaves in plants grown in Full or LowN treatments. Data collected every 5 days from 10 DAE. Points show the coefficient estimated for each treatment, with bars extending +/-1 standard error (SE). Colored polygons follow SE bars. B) as A) for total leaf area (TLA). Endpoint coefficients (+/-1 SE) for root occupancy (root fresh weight/total plant fresh weight as a percentage) in C) the upper 45 cm (R_0-45_) of the soil and D) below 45 cm (R_45+_) in the soil, for LowN or LowP treatments. Significant pairwise differences in Tukey HSD test (*α* = 0.1) indicated by lowercase letters.

We used inductively coupled plasma mass spectrometry to quantify total P and 19 further elements in leaf tissue harvested from the segregating mutant stock (Supplemental Data Set 1E). There was no significant effect of *pho1;2a* on total P concentration in the leaf. A previously reported increase in leaf P concentration under lowN (Schlüter et al., 2013; Torres-Rodríguez et al., 2021) was observed in all genotypes, further indicating that *pho1;2a* mutants are not compromised in the translocation of P from roots to leaves. Although we saw treatment effects on several elements, differences between genotypes were subtle, with limited statistical support that was not robust to correction for multiple testing (Supplemental Figure 3; Supplemental Data Set 1E).

### Root density distribution in *pho1;2a* mutation is adjusted in response to low nutrient availability

Root biomass showed a mild reduction in homozygous mutants and heterozygotes compared with wild-type plants, again with the greatest statistical support under LowN (Supplemental Figure 2 - RFW LowN; Supplemental Data Set 1D). In the same way, the maximum number of roots is decreased in homozygous mutants under LowN (Supplemental Figure 4 - MaxNR LowN; Supplemental Data Set 1F). There was a reduction in the mean number of crown roots in mutants specifically under LowP (wt: 15.02 ± 1.15, ht: 14.21 ± 0.93, mt: 12.14 ± 1.28; here and below, we give model coefficients and associated standard errors, Supplemental Figure 2 - CN LowP), although without strong statistical support. Mutants showed a significant reduction in root growth in the upper 45 cm of the soil column under LowN, in terms of both absolute biomass and proportion of total biomass (Fig. 2C; Supplemental Figure 2; Supplemental Data set 1D). A similar, although non-significant, trend was also seen under LowP (Fig. 2C). In contrast to the shoot growth traits, the action of the *pho1;2a* mutation on root biomass appeared additive, with heterozygous plants intermediate to homozygous mutants and wild-type. Interestingly, in LowN and LowP, homozygous mutant and heterozygous plants showed greater root occupancy in the lower 45 cm of the soil column than wild-type, both in terms of absolute mean biomass and mean proportion of total biomass (Fig. 2D). The trend towards deeper rooting in mutants was reversed under LowNP, with homozygous mutant and heterozygous plants showing greater biomass than wild-type in the top 45 cm and less than wild-type in the lower 45 cm, the latter difference being detected as significant in both absolute and proportional terms.

### Root and leaf transcriptomes are modified in *pho1;2a* mutants

The *Arabidopsis Pho1* gene is involved not only in Pi translocation but also in nutrient signaling (Rouached et al., 2011). Although maize *pho1;2a* mutants lacked a strong Pi translocation phenotype, we speculated that subtle growth differences in the mutant might result from a signaling effect. To evaluate nutritional signaling in *pho1;2a* mutants, we grew a second batch of segregating plants under the same treatments used for growth evaluation. At 25 DAE (the point at which the nutrient treatments first had a clear significant effect on growth), we harvested leaf and root tissue from homozygous mutants, heterozygotes and wild-type plants for transcriptome analysis. To identify *Pho1;2a* regulated genes in our transcriptome data, we first compared genotypes within any given treatment/tissue combination and then pooled information to obtain statistical support for genotype differences, considering wild-type to heterozygote (wt-ht) and wild-type to homozygous mutant (wt-mt) comparisons separately.

We identified 1,100 *Pho1;2a*-regulated genes combining the wt-ht and wt-mt comparisons, the majority upregulated in *pho1;2a* mutants or heterozygotes with respect to wild-type (Fig. 3; Supplemental Data Set 2A; Supplemental Figure 5). We saw a greater number of expression differences in roots than leaves, consistent with the expression pattern of *Pho1;2a*. The number of expression differences in roots was similar under all four nutrient treatments (Fig. 3A). In addition, the majority of genes were shared across at least two treatments, with a high proportion shared across all treatments (Fig. 3B). In leaves, the great majority of expression differences were seen under LowP or LowNP (Fig. 3A, C). A smaller proportion of leaf regulated genes was shared across two or more treatments than in roots (Fig. 3C). The number of expression differences across treatment/tissue combinations was similar in wt-ht and wt-mt comparisons, indicating a partially dominant effect of *pho1;2a* on transcript accumulation (Supplemental Figure 5).

**Figure 3.**
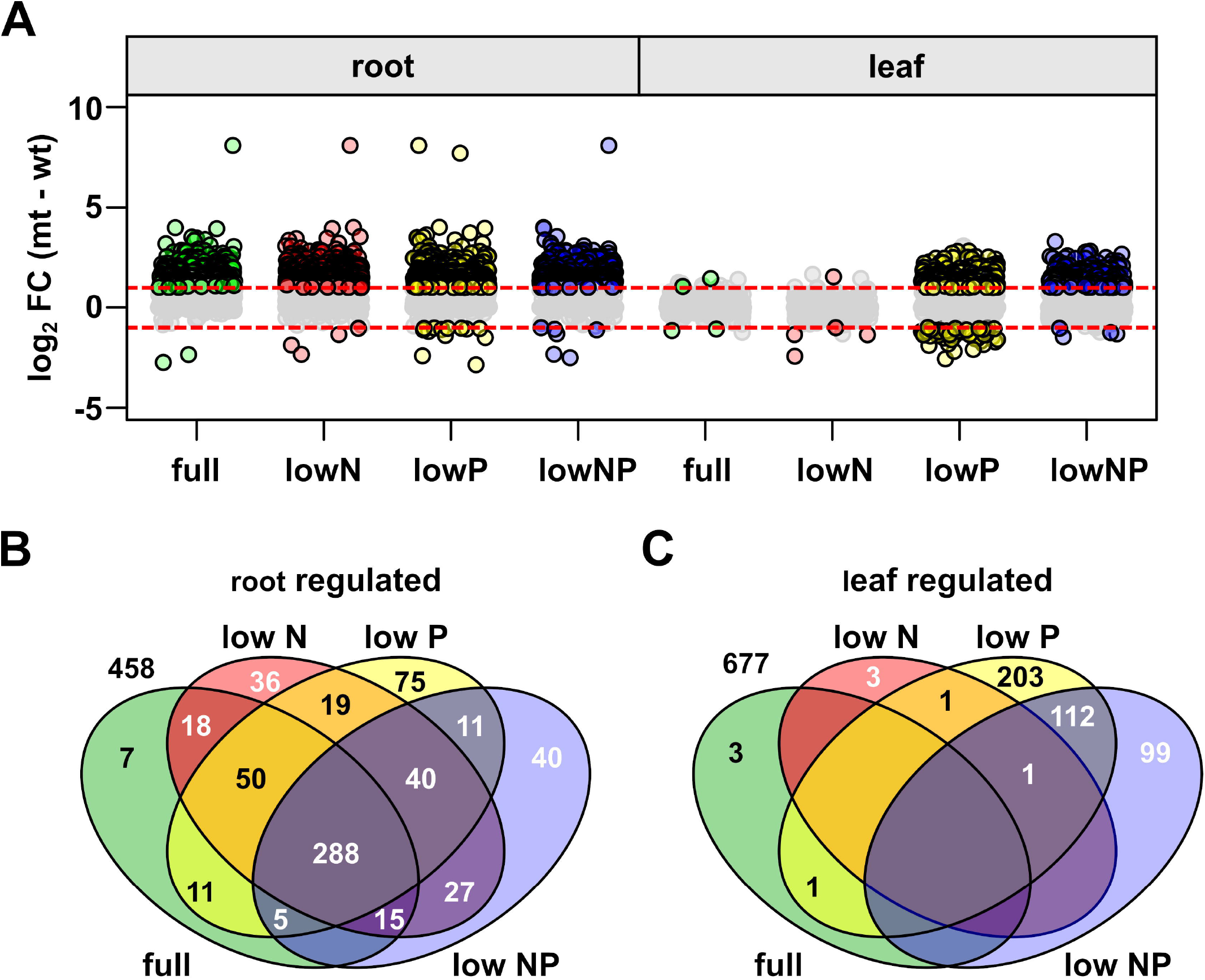
The *pho1;2a’m1.1* mutation is linked to transcriptional misregulation. A) Effect size estimates (log_2_ fold change; homozygous - wild-type) for genes differentially expressed in plants homozygous for *pho1;2a* compared with segregating wild-type siblings. Points corresponding to significant genes in any given tissue-treatment combination are filled and outlined in bold. A single gene with an effect size >5 in the roots is not shown. B) Venn diagram showing the overlap (count) of genes differentially expressed in the root (mutant - wild-type) among the four nutrient treatments. Genes from the 1,100 list not belonging to any set are not shown. C) as B, for genes differentially expressed in the leaf.

To investigate an effect on P signaling, we considered the response of *Pho1;2a* regulated genes to LowP in wild-type and mutant plants (Supplemental Data Set 2B, C). A total of 581 of the 1,100-gene set were called as *Pho1;2a-*regulated in LowP roots (combining wt-mt and mt-ht comparisons). Overall, these 581 genes showed a similar pattern of root expression under Full nutrients, genotype differences being maintained, but with no evidence of a treatment effect (Fig. 4A). In contrast, the 366 of the 1,100-gene set that were called as *Pho1;2a-*regulated in LowP leaves, were broadly not differentially expressed among genotypes under Full nutrients, but did show evidence of induction under LowP - *i*.*e*. this group of 366 genes were induced by LowP in wild-type plants, but induced to a greater extent in plants carrying the *pho1;2a* mutation (Fig. 4B). We interpret this pattern as evidence of mis-regulation of the P starvation response in the leaves of *pho1;2a* mutants. The *Pho1;2a* regulated gene set was enriched (Supplemental Data Set 2D) for GO terms associated with the P starvation including the specific term GO16036: cellular response to P starvation itself (PS) and several terms related to phytohormone signaling. Genes associated with abiotic stress GO terms followed the global trend of upregulation in mutant roots across all treatments, and upregulation specifically under LowP and LowNP in leaves (*e*.*g*. GO:9737 response to abscisic acid, GO: 9753 response to jasmonic acid, GO: 10167 response to nitrate. Fig. 4C-E). For the P starvation term, upregulation in mutants was LowP specific in both roots and leaves (Fig. 4F). Overall, we interpret the transcriptome data to implicate *Pho1;2a* in signaling. Specifically, we highlight LowP-dependent expression differences in mutant leaves as an indication of a role in systemic P signaling.

**Figure 4.**
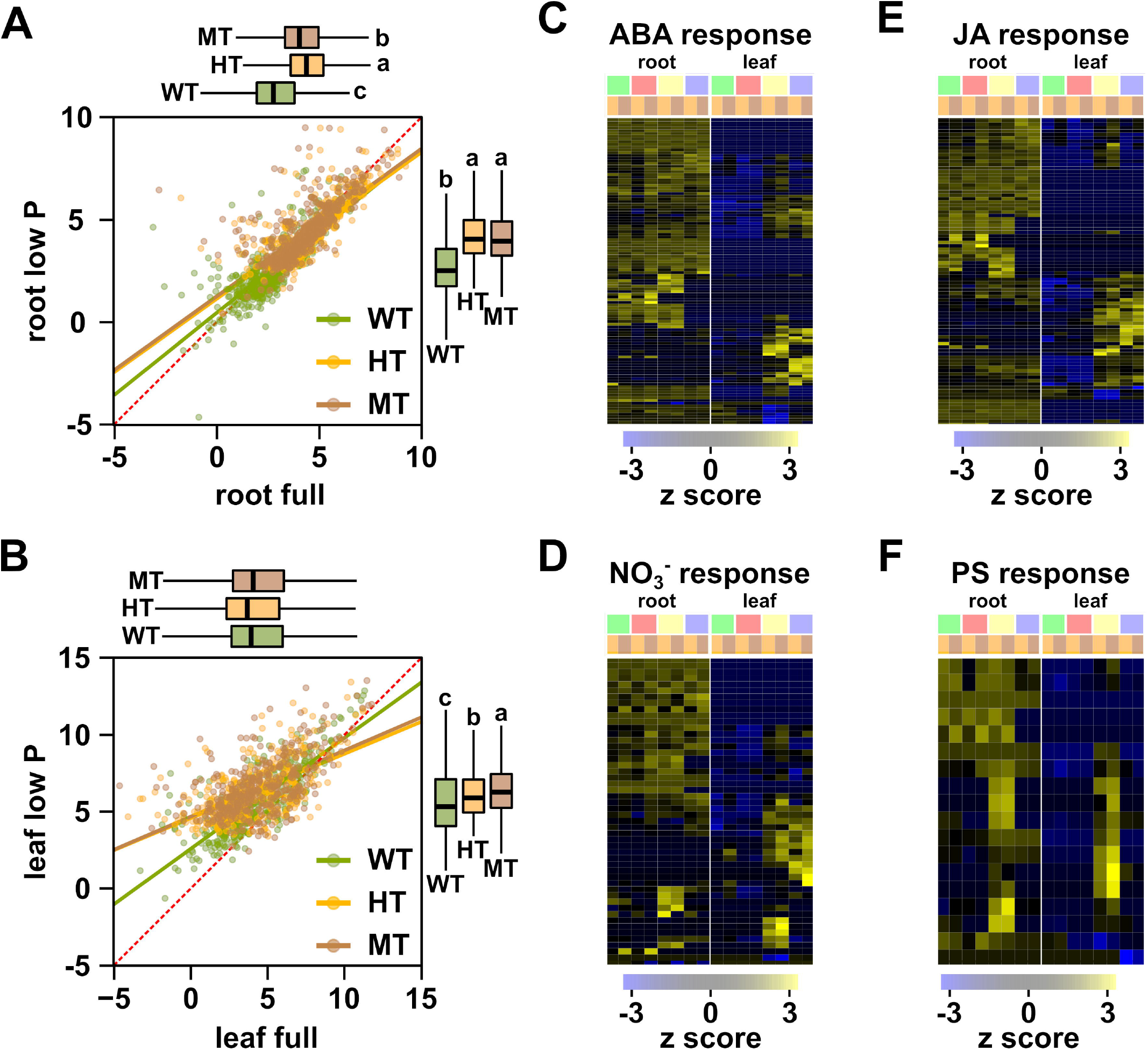
The phosphate starvation response is promoted in the leaves of *pho1;2a* mutants. A) Scatter plot of root expression coefficients (∼standardized expression) comparing control (root full) and LowP (root low P) for a set of 581 genes identified as *Pho1;2a-*regulated in LowP roots. Expression shown for homozygous (MT), heterozygous (HT) or wild type (WT) plants. A linear fit for each genotype is shown by the colored solid lines. The dashed red line marks equal expression in the two treatments. Marginal box plots indicate the distribution of values with respect to the facing axis. Where there was a significant genotype effect (Kruskal-Wallis test, p < 0.001), lower case letters indicate groups defined by Dunn’s test (p < 0.001). B) as A, showing a set of 366 genes identified as *Pho1;2a-*regulated in lowP leaves. C) Effect size estimates (standardized by row; z-score) of the *pho1;2a* mutation on expression of called genes associated with abscisic acid (ABA) response (GO:9737). Columns represent conditions (root or leaf as stated; treatments indicated as green bar: Full, red: LowN, yellow: LowP, blue: LowNP; genotype as orange: heterozygous - wild type, brown: homozygous - wild type). Each row is a different gene. D - F) as C, for genes associated with jasmonic acid (JA; GO: 9753), nitrate (NO_3_^-^; GO: 10167) and cellular response to phosphorus starvation (PS; GO: 16036) responses, respectively.

## DISCUSSION

We have shown that insertion of 8 bp in the 6th exon of the maize *Pho1;2a* gene is correlated with mild growth-reduction and mis-regulation of the transcriptional response to low P availability. In *Arabidopsis*, the PHO1 protein functions in the translocation of Pi from roots to shoots and in P signaling (Hamburger et al., 2002; Secco et al., 2010; Rouached et al., 2011). As a consequence, loss-of-function *pho1* mutants in *Arabidopsis* show a dramatic reduction in shoot P concentration and growth (Poirier et al., 1991). A similar, non-redundant role in P translocation is played by the orthologous proteins PHO1;2 and PHO1;1 in rice and tomato (Secco et al., 2010; Zhao et al., 2019), respectively, although shoot growth can recover by maturity in rice (Ma et al., 2021). The single copy rice *PHO1;2* is represented by the paralogous gene pair *Pho1;2a* and *Pho1;2b* in maize (Salazar-Vidal et al., 2016). Characterization of the *pho1;2a* allele presented here indicates that the maize PHO1;2a protein does not contribute significantly to Pi translocation. There was no significant reduction in leaf P concentration in *pho1;2a* mutants nor any severe impact of growth, such as has been seen in other plant species. We observed *Pho1;2b* transcripts in the root to accumulate to levels ∼40-fold greater than those of *Pho1;2a*, consistent with *Pho1;2b* playing the primary role in P translocation. We saw no evidence of transcriptional up-regulation of *Pho1;2b* in *pho1;2a* mutants, suggesting the absence of any masking of a loss-of-function phenotype by the compensatory action of the paralogous gene. In mature plants, *Pho1;2a* and *Pho1;2b* have been shown to play non-redundant roles in grain filling (Ma et al., 2021), although their expression levels are more nearly equivalent in the endosperm (Zhan et al., 2015). Interestingly, the previous study did not report any dramatic growth phenotype associated with either *pho1;2a* or *pho1;2b* single mutants, suggesting that double or higher-order mutations in the maize *pho1* family might be required to produce the P translocation phenotype seen in other plants.

We have interpreted our observations under the assumption that the *pho1;2a’-m1*.*1* allele conditions a complete loss of PHO1;2a protein function. Of course, this would be better demonstrated by isolation of additional mutant alleles. Similarly, the level of redundancy between the *Pho1;2a* and *Pho1;2b* paralogs cannot really be addressed without generation of *pho1;2b* mutants. Although we are actively pursuing a number of avenues to obtain such mutants, no such material is currently available. The CRISPR/Cas9 edit reported by Ma et al. (Ma et al., 2021) is clearly an important additional reference, although the molecular nature of the Ma et al. *pho1;2a* allele - a small deletion in Exon 6 - is very similar to the *pho1;2a’-m1*.*1* footprint event. PHO1 proteins contain both an N-terminal SPX domain and a C-terminal EXS domain. The molecular lesions in both *pho1;2a’-m1*.*1* and the Ma et al. *pho1;2a* allele are located downstream of the SPX domain-encoding portion of the gene and, as a consequence, have the potential to generate truncated SPX-containing proteins. In *Arabidopsis*, the PHO1 EXS domain is required for subcellular localization, Pi transport and root-to-shoot signaling (Wang et al., 2004; Stefanovic et al., 2007; Secco et al. 2012; Wege et al., 2016). In the absence of the EXS domain, the function of a truncated maize *pho1;2a* product would clearly be greatly compromised. Nevertheless, there is an entire class of single SPX domain proteins that play an important role in P signaling mediated through protein-protein interaction (Poirier, 2019; Torres-Rodríguez et al., 2021; Xiao et al., 2021). Experimental manipulation has demonstrated the stability and nuclear localization of an isolated PHO1 SPX domain in *Arabidopsis* (Wege et al., 2016). Furthermore, overexpression of the SPX domain of *Arabidopsis* PHO1;H4 is sufficient to promote constitutive hypocotyl elongation, mimicking the dominant *Pho1;H4 shb1-D* mutant (Zhou and Ni, 2010). It may be significant that transcriptional effects associated with the maize *pho1;2a’-m1*.*1* mutant were mostly dominant, although our observations would also be consistent with previously reported *PHO1* dosage-effects (Hamburger et al., 2002; Secco et al., 2010; Rouached et al., 2011). Ma et al. do not comment on the dominance of kernel phenotypes associated with *pho1;2a* and *pho1;2b* CRISPR/Cas-9 edits (Ma et al., 2021). As for redundancy, isolation of additional mutant alleles will clarify the interpretation of dominance and dosage.

The presence of both *Pho1;2a* and *Pho1;2b* in the maize genome, coupled with their syntenic relationship to the single *Pho1;2* gene of the closely related plant sorghum, suggests that the two maize genes are a paralog pair retained following a lineage specific whole genome duplication (Salazar-Vidal et al., 2016). For the majority of genes, fractionation following whole-genome duplication restores gene copy number to pre-duplication levels (Schnable et al., 2011). The retention of *Pho1;2a*, despite a low level of expression relative to *Pho1;2b* in most tissues analyzed, suggests the gene to have functional significance, although the fractionation process may be ongoing. In addition to the reported role in grain filling (Ma et al., 2021), it is interesting to note that *Pho1;2a* has been identified as a candidate in several quantitative trait loci mapping and genome wide association studies, including studies of variation in P efficiency (Zhang et al., 2014) and root system architecture (Wang et al., 2019). Furthermore, *Pho1;2a* is located within a large-scale inversion polymorphism that is characteristic of the native maize varieties and maize wild-relatives of the central Mexican highlands (Crow et al., 2020) and has been associated with low temperature and soil P availability (Aguirre-Liguori et al., 2019). To our knowledge, *Pho1;2b* has not been associated with quantitative trait variation in maize. It may be significant that *Pho1;2a* has no major role in Pi translocation, freeing it up for possible sub- or neo-functionalization and a role in adaptive fine-tuning to different environments.

In *Arabidopsis, PHO1* plays a role in both Pi translocation and systemic P signaling (Rouached et al., 2011; Wege et al., 2016). We found evidence for transcriptional upregulation of several hundred genes in *pho1;2a* mutant and heterozygous plants. In roots, increased transcript accumulation was seen under all nutrient regimes, the majority of upregulated genes being shared across treatments. In contrast, we only saw a substantial effect in leaves when plants were grown under LowP or LowNP conditions. Maize *Pho1;2a* transcripts were only detected in roots. Similarly, *Arabidopsis PHO1* and rice *Pho1;2* are preferentially expressed in root cells of the stele, including the pericycle and xylem parenchyma cells (Hamburger et al., 2002; Jabnoune et al., 2013). We saw many more significantly upregulated transcripts than we did down regulated transcripts, suggesting a primarily repressive role for *Pho1;2a*. Interestingly, many of the transcripts and associated GO terms upregulated in *pho1;2a* mutants have previously been found to be upregulated under low P in wild-type plants (Torres-Rodríguez et al., 2021). In the case of the leaves, transcripts accumulating to higher levels in *pho1;2a* mutant and heterozygous plants under low P availability were at higher levels in all genotypes than seen under full conditions. Several transcripts directly assigned to the P starvation GO term (GO: 16036) were upregulated in both roots and leaves under LowP, the level of induction being higher in *pho1;2a* mutants than in wild-type plants (Fig. 4F). We interpret these results to suggest that the transcriptional P starvation response is exaggerated in *pho1;2a* mutants and heterozygous plants. It would be informative to characterize the dynamics of the induction of the P starvation response in *pho1;2a* mutants to assess a possible role in fine-tuning or feedback regulation.

Transcripts associated with several phytohormone GO terms were upregulated in *pho1;2a* mutant and heterozygous plants. Induction of phytohormone signaling is consistent with a constitutive or exaggerated P starvation response. *Arabidopsis* PHO1 has been shown previously to be involved in ABA mediated germination (Huang et al., 2017) and stomatal opening responses (Zimmerli et al., 2012). Furthermore, in *Arabidopsis* the level of induction of the jasmonic acid regulated genes *JAZ10, VSP2* and *LOX2* following wounding is greater in *pho1* mutants than in wild-type plants (Khan et al., 2016), showing a pattern analogous to that which we observed in maize *pho1;2a* mutants under low P availability.

In summary, we show that full PHO1;2a is not required for root to shoot P translocation in maize, although the *pho1;2a’-m1*.*1* triggers transcriptional mis-regulation indicating a role in signaling. In the future, it will be informative to evaluate the role of the paralogous *Pho1;2b* gene, and indeed the further two members of the maize *Pho1* gene family. Furthermore, characterization of *pho1;2a; pho1;2b* double mutants will be necessary to better understand functional divergence and redundancy between the two maize *Pho1;2* genes.

## Supporting information

Supplemental Data Set 1

Supplemental Data Set 2

## ACKNOWLEDGEMENTS

We thank Arturo Chavez-Ramirez, Dario Alavez-Mercado, Oscar Nieves, Benjamin Barrales-Gamez, and Jessica Carcaño-Macias and for technical assistance in plant measurement and tissue collection. We thank Rocio Martinez for her enthusiastic help and support in greenhouse conditions. We thank Labsergen (Laboratorio de Servicios Genómicos, Langebio) for guidance and assistance in the preparation and sequencing of RNA libraries. We thank Ricardo Chávez Montes, Daniel Runcie and Taylor Crow for invaluable help with analysis of RNA sequence data. We thank Ivan Baxter and Greg Ziegler for supporting ICP-MS analysis.

This work was supported by the Mexican Consejo Nacional de Ciencia y Tecnología (CB-2015-01 254012). ALAN is supported by the CONACyT graduate fellowship number 486795. RJHS is supported by the USDA National Institute of Food and Agriculture and Hatch Appropriations under Project #PEN04734 and Accession #1021929. RNA Sequencing at UC Berkeley was partially supported through NIH S10 OD018174 Instrumentation Grant.

## LITERATURE CITED

Aguirre-Liguori JA, Gaut BS, Jaramillo-Correa JP, Tenaillon MI, Montes-Hernández S, García-Oliva F, Hearne SJ, Eguiarte LE (2019) Divergence with gene flow is driven by local adaptation to temperature and soil phosphorus concentration in teosinte subspecies (Zea mays parviglumis and Zea mays mexicana). Mol Ecol 28: 2814–2830

Ai P, Sun S, Zhao J, Fan X, Xin W, Guo Q, Yu L, Shen Q, Wu P, Miller AJ, et al (2009) Two rice phosphate transporters, OsPht1;2 and OsPht1;6, have different functions and kinetic properties in uptake and translocation. The Plant Journal 57: 798–809

Bai L, Singh M, Pitt L, Sweeney M, Brutnell TP (2007) Generating novel allelic variation through Activator insertional mutagenesis in maize. Genetics 175: 981–992

Baldwin T, Sakthianandeswaren A, Curtis JM, Kumar B, Smyth GK, Foote SJ, Handman E (2007) Wound healing response is a major contributor to the severity of cutaneous leishmaniasis in the ear model of infection. Parasite Immunology 29: 501–513

Barragán-Rosillo AC, Peralta-Alvarez CA, Ojeda-Rivera JO, Arzate-Mejía RG, Recillas-Targa F, Herrera-Estrella L (2021) Genome accessibility dynamics in response to phosphate limitation is controlled by the PHR1 family of transcription factors in Arabidopsis. Proc Natl Acad Sci U S A. DOI: 10.1073/pnas.2107558118

Baxter IR, Vitek O, Lahner B, Muthukumar B, Borghi M, Morrissey J, Guerinot ML, Salt DE (2008) The leaf ionome as a multivariable system to detect a plant’s physiological status. Proc Natl Acad Sci U S A 105: 12081–12086

Bray NL, Pimentel H, Melsted P, Pachter L (2016) Near-optimal probabilistic RNA-seq quantification. Nat Biotechnol 34: 525–527

Bulak Arpat A, Magliano P, Wege S, Rouached H, Stefanovic A, Poirier Y (2012) Functional expression of PHO1 to the Golgi and trans-Golgi network and its role in export of inorganic phosphate. Plant J 71: 479–491

Bustos R, Castrillo G, Linhares F, Puga MI, Rubio V, Pérez-Pérez J, Solano R, Leyva A, Paz-Ares J (2010) A central regulatory system largely controls transcriptional activation and repression responses to phosphate starvation in Arabidopsis. PLoS Genet 6: e1001102

Chang MX, Gu M, Xia YW, Dai XL, Dai CR, Zhang J, Wang SC, Qu HY, Yamaji N, Feng Ma J, et al (2019) OsPHT1;3 mediates uptake, translocation, and remobilization of phosphate under extremely low phosphate regimes. Plant Physiol 179: 656–670

Chien P-S, Chiang C-P, Leong SJ, Chiou T-J (2018) Sensing and signaling of phosphate starvation: from local to long distance. Plant Cell Physiol 59: 1714–1722

Crow T, Ta J, Nojoomi S, Aguilar-Rangel MR, Torres Rodríguez JV, Gates D, Rellán-Álvarez R, Sawers R, Runcie D (2020) Gene regulatory effects of a large chromosomal inversion in highland maize. PLoS Genet 16: e1009213

Diepenbrock W (1991) Properties of root membrane lipids as related to mineral nutrition. In BL McMichael, H Persson, eds, Developments in Agricultural and Managed Forest Ecology. Elsevier, pp 25–30

Escobar MA, Geisler DA, Rasmusson AG (2006) Reorganization of the alternative pathways of the Arabidopsis respiratory chain by nitrogen supply: opposing effects of ammonium and nitrate. Plant J 45: 775–788

Francis CA, Rutger JN, Palmer AFE (1969) A rapid method for plant leaf area estimation in maize (Zea mays L.) 1. Crop Sci 9: 537–539

Galkovskyi T, Mileyko Y, Bucksch A, Moore B, Symonova O, Price CA, Topp CN, Iyer-Pascuzzi AS, Zurek PR, Fang S, et al (2012) GiA Roots: software for the high throughput analysis of plant root system architecture. BMC Plant Biol 12: 116

Gonzalez-Segovia E, Pérez-Limon S, Cíntora-Martínez GC, Guerrero-Zavala A, Janzen GM, Hufford MB, Ross-Ibarra J, Sawers RJH (2019) Characterization of introgression from the teosinte Zea mays ssp. mexicana to Mexican highland maize. PeerJ 7: e6815

Hamburger D, Rezzonico E, MacDonald-Comber Petétot J, Somerville C, Poirier Y (2002) Identification and characterization of the Arabidopsis PHO1 gene involved in phosphate loading to the xylem. Plant Cell 14: 889–902

Hinsinger P (2001) Bioavailability of soil inorganic P in the rhizosphere as affected by root-induced chemical changes: a review. Plant and Soil 237: 173–195

Hoagland DR, Broyer TC (1936) General nature of the process of salt accumulation by roots with description of experimental methods. Plant Physiol 11: 471–507

Huang Y, Sun M-M, Ye Q, Wu X-Q, Wu W-H, Chen Y-F (2017) Abscisic acid modulates seed germination via ABA INSENSITIVE5-Mediated PHOSPHATE1. Plant Physiol 175: 1661–1668

Hu B, Jiang Z, Wang W, Qiu Y, Zhang Z, Liu Y, Li A, Gao X, Liu L, Qian Y, et al (2019) Nitrate–NRT1.1B–SPX4 cascade integrates nitrogen and phosphorus signalling networks in plants. Nature Plants 5: 401–413

Jabnoune M, Secco D, Lecampion C, Robaglia C, Shu Q, Poirier Y (2013) A rice cis-natural antisense RNA acts as a translational enhancer for its cognate mRNA and contributes to phosphate homeostasis and plant fitness. Plant Cell 25: 4166–4182

Khan GA, Vogiatzaki E, Glauser G, Poirier Y (2016) Phosphate deficiency induces the jasmonate pathway and enhances resistance to insect herbivory. Plant Physiol 171: 632–644

Lin F, Jiang L, Liu Y, Lv Y, Dai H, Zhao H (2014) Genome-wide identification of housekeeping genes in maize. Plant Mol Biol 86: 543–554

Lin W-Y, Huang T-K, Leong SJ, Chiou T-J (2014) Long-distance call from phosphate: systemic regulation of phosphate starvation responses. J Exp Bot 65: 1817–1827

López-Arredondo DL, Leyva-González MA, González-Morales SI, López-Bucio J, Herrera-Estrella L (2014) Phosphate nutrition: improving low-phosphate tolerance in crops. Annu Rev Plant Biol 65: 95–123

Lynch J, Epstein E, Lauchli A, Weight GI (1990) An automated greenhouse sand culture system suitable for studies of P nutrition. Plant Cell Environ 13: 547–554

Ma B, Zhang L, Gao Q, Wang J, Li X, Wang H, Liu Y, Lin H, Liu J, Wang X, et al (2021) A plasma membrane transporter coordinates phosphate reallocation and grain filling in cereals. Nat Genet. DOI: 10.1038/s41588-021-00855-6

Maere S, Heymans K, Kuiper M (2005) BiNGO: a Cytoscape plugin to assess overrepresentation of gene ontology categories in biological networks. Bioinformatics 21: 3448–3449

McCarthy DJ, Chen Y, Smyth GK (2012) Differential expression analysis of multifactor RNA-Seq experiments with respect to biological variation. Nucleic Acids Res 40: 4288–4297

de Mendiburu F (2020) agricolae: Statistical Procedures for Agricultural Research.

Misson J, Thibaud M-C, Bechtold N, Raghothama K, Nussaume L (2004) Transcriptional regulation and functional properties of Arabidopsis Pht1;4, a high affinity transporter contributing greatly to phosphate uptake in phosphate deprived plants. Plant Mol Biol 55: 727–741

Paz-Ares J, Puga MI, Rojas-Triana M, Martinez-Hevia I, Diaz S, Poza-Carrión C, Miñambres M, Leyva A (2022) Plant adaptation to low phosphorus availability: core signaling, crosstalks, and applied implications. Mol Plant 15: 104–124

Péret B, Clément M, Nussaume L, Desnos T (2011) Root developmental adaptation to phosphate starvation: better safe than sorry. Trends Plant Sci 16: 442–450

Plaxton WC, Tran HT (2011) Metabolic adaptations of phosphate-starved plants. Plant Physiol 156: 1006–1015

Poirier Y (2019) Post-translational Regulation of SPX Proteins for coordinated nutrient signaling. Mol Plant 12: 1041–1043

Poirier Y, Thoma S, Somerville C, Schiefelbein J (1991) Mutant of Arabidopsis deficient in xylem loading of phosphate. Plant Physiol 97: 1087–1093

Puga MI, Mateos I, Charukesi R, Wang Z, Franco-Zorrilla JM, de Lorenzo L, Irigoyen ML, Masiero S, Bustos R, Rodríguez J, et al (2014) SPX1 is a phosphate-dependent inhibitor of Phosphate Starvation Response 1 in Arabidopsis. Proc Natl Acad Sci U S A 111: 14947–14952

Raghothama KG (1999) Phosphate acquisition. Annu Rev Plant Physiol Plant Mol Biol 50: 665–693

Ramírez-Flores MR, Rellán-Álvarez R, Wozniak B, Gebreselassie M-N, Jakobsen I, Olalde-Portugal V, Baxter I, Paszkowski U, Sawers RJH (2017) Co-ordinated changes in the accumulation of metal ions in maize (Zea mays ssp. mays L.) in response to inoculation with the arbuscular mycorrhizal fungus Funneliformis mosseae. Plant Cell Physiol 58: 1689–1699

Rausch C, Bucher M (2002) Molecular mechanisms of phosphate transport in plants. Planta 216: 23–37

R Core Team (2022) R: A Language and Environment for Statistical Computing.

Reddy AR, Reddy KR, Padjung R, Hodges HF (1996) Nitrogen nutrition and photosynthesis in leaves of Pima cotton. J Plant Nutr 19: 755–770

Reis RS, Deforges J, Schmidt RR, Schippers JHM, Poirier Y (2021) An antisense noncoding RNA enhances translation via localized structural rearrangements of its cognate mRNA. Plant Cell 33: 1381–1397

Ried MK, Wild R, Zhu J, Pipercevic J, Sturm K, Broger L, Harmel RK, Abriata LA, Hothorn LA, Fiedler D, et al (2021) Inositol pyrophosphates promote the interaction of SPX domains with the coiled-coil motif of PHR transcription factors to regulate plant phosphate homeostasis. Nat Commun 12: 384

Ritchie ME, Phipson B, Wu D, Hu Y, Law CW, Shi W, Smyth GK (2015) limma powers differential expression analyses for RNA-sequencing and microarray studies. Nucleic Acids Research 43: e47

Robinson MD, McCarthy DJ, Smyth GK (2010) edgeR: a Bioconductor package for differential expression analysis of digital gene expression data. Bioinformatics 26: 139–140

Rouached H, Stefanovic A, Secco D, Bulak Arpat A, Gout E, Bligny R, Poirier Y (2011) Uncoupling phosphate deficiency from its major effects on growth and transcriptome via PHO1 expression in Arabidopsis. Plant J 65: 557–570

Salazar-Vidal MN, Acosta-Segovia E, Sánchez-León N, Ahern KR, Brutnell TP, Sawers RJH (2016) Characterization and transposon mutagenesis of the maize (Zea mays) Pho1 gene family. PLoS One 11: e0161882

Schachtman DP, Reid RJ, Ayling SM (1998) Phosphorus uptake by plants: from soil to cell. Plant Physiol 116: 447–453

Schlüter U, Colmsee C, Scholz U, Bräutigam A, Weber APM, Zellerhoff N, Bucher M, Fahnenstich H, Sonnewald U (2013) Adaptation of maize source leaf metabolism to stress related disturbances in carbon, nitrogen and phosphorus balance. BMC Genomics 14: 442

Schnable JC, Springer NM, Freeling M (2011) Differentiation of the maize subgenomes by genome dominance and both ancient and ongoing gene loss. Proc Natl Acad Sci U S A 108: 4069–4074

Secco D, Baumann A, Poirier Y (2010) Characterization of the rice PHO1 gene family reveals a key role for OsPHO1;2 in phosphate homeostasis and the evolution of a distinct clade in dicotyledons. Plant Physiol 152: 1693–1704

Secco D, Wang C, Arpat BA, Wang Z, Poirier Y, Tyerman SD, Wu P, Shou H, Whelan J (2012) The emerging importance of the SPX domain-containing proteins in phosphate homeostasis. New Phytol 193: 842–851

Shin H, Shin H-S, Dewbre GR, Harrison MJ (2004) Phosphate transport in Arabidopsis: Pht1;1 and Pht1;4 play a major role in phosphate acquisition from both low-and high-phosphate environments. Plant J 39: 629–642

Soneson C, Love MI, Robinson MD (2015) Differential analyses for RNA-seq: transcript-level estimates improve gene-level inferences. F1000Res 4: 1521

Stefanovic A, Arpat AB, Bligny R, Gout E, Vidoudez C, Bensimon M, Poirier Y (2011) Over-expression of PHO1 in Arabidopsis leaves reveals its role in mediating phosphate efflux. Plant J 66: 689–699

Stefanovic A, Ribot C, Rouached H, Wang Y, Chong J, Belbahri L, Delessert S, Poirier Y (2007) Members of the PHO1 gene family show limited functional redundancy in phosphate transfer to the shoot, and are regulated by phosphate deficiency via distinct pathways. Plant J 50: 982–994

Thibaud M-C, Arrighi J-F, Bayle V, Chiarenza S, Creff A, Bustos R, Paz-Ares J, Poirier Y, Nussaume L (2010) Dissection of local and systemic transcriptional responses to phosphate starvation in Arabidopsis. Plant J 64: 775–789

Torres-Rodríguez JV, Salazar-Vidal MN, Chávez Montes RA, Massange-Sánchez JA, Gillmor CS, Sawers RJH (2021) Low nitrogen availability inhibits the phosphorus starvation response in maize (Zea mays ssp. mays L.). BMC Plant Biol 21: 259

Ueda Y, Kiba T, Yanagisawa S (2020) Nitrate-inducible NIGT1 proteins modulate phosphate uptake and starvation signalling via transcriptional regulation of SPX genes. Plant J 102: 448–466

Urbut S, Wang G, Carbonetto P, Stephens M (2019) Flexible statistical methods for estimating and testing effects in genomic studies with multiple conditions. Nature Genetics 51: 187–195

Vance CP, Uhde-Stone C, Allan DL (2003) Phosphorus acquisition and use: critical adaptations by plants for securing a nonrenewable resource. New Phytol 157: 423–447

Veneklaas EJ, Lambers H, Bragg J, Finnegan PM, Lovelock CE, Plaxton WC, Price CA, Scheible W-R, Shane MW, White PJ, et al (2012) Opportunities for improving phosphorus-use efficiency in crop plants. New Phytol 195: 306–320

Wang Q-J, Yuan Y, Liao Z, Jiang Y, Wang Q, Zhang L, Gao S, Wu F, Li M, Xie W, et al (2019) Genome-wide association study of 13 traits in maize seedlings under low phosphorus stress. Plant Genome 12: 190039

Wang Y, Ribot C, Rezzonico E, Poirier Y (2004) Structure and expression profile of the Arabidopsis PHO1 gene family indicates a broad role in inorganic phosphate homeostasis. Plant Physiol 135: 400–411

Wang Z, Ruan W, Shi J, Zhang L, Xiang D, Yang C, Li C, Wu Z, Liu Y, Yu Y, et al (2014) Rice SPX1 and SPX2 inhibit phosphate starvation responses through interacting with PHR2 in a phosphate-dependent manner. Proc Natl Acad Sci U S A 111: 14953–14958

Wege S, Khan GA, Jung J-Y, Vogiatzaki E, Pradervand S, Aller I, Meyer AJ, Poirier Y (2016) The EXS domain of PHO1 participates in the response of shoots to phosphate deficiency via a root-to-shoot signal. Plant Physiol 170: 385–400

Woodhouse MR, Sen S, Schott D, Portwood JL, Freeling M, Walley JW, Andorf CM, Schnable JC (2021) qTeller: A tool for comparative multi-genomic gene expression analysis. Bioinformatics. DOI: 10.1093/bioinformatics/btab604

Xiao J, Xie X, Li C, Xing G, Cheng K, Li H, Liu N, Tan J, Zheng W (2021) Identification of SPX family genes in the maize genome and their expression under different phosphate regimes. Plant Physiol Biochem 168: 211–220

Zhang H, Uddin MS, Zou C, Xie C, Xu Y, Li W-X (2014) Meta-analysis and candidate gene mining of low-phosphorus tolerance in maize. J Integr Plant Biol 56: 262–270

Zhan J, Thakare D, Ma C, Lloyd A, Nixon NM, Arakaki AM, Burnett WJ, Logan KO, Wang D, Wang X, et al (2015) RNA sequencing of laser-capture microdissected compartments of the maize kernel identifies regulatory modules associated with endosperm cell differentiation. Plant Cell 27: 513–531

Zhao P, You Q, Lei M (2019) A CRISPR/Cas9 deletion into the phosphate transporter SlPHO1;1 reveals its role in phosphate nutrition of tomato seedlings. Physiol Plant 167: 556–563

Zhou Y, Ni M (2010) SHORT HYPOCOTYL UNDER BLUE1 truncations and mutations alter its association with a signaling protein complex in Arabidopsis. Plant Cell 22: 703–715

Zimmerli C, Ribot C, Vavasseur A, Bauer H, Hedrich R, Poirier Y (2012) PHO1 expression in guard cells mediates the stomatal response to abscisic acid in Arabidopsis. Plant J 72: 199–211

