## Supplementary figures and images for "The *pho1;2a’-m1.1* allele of *Phosphate1* conditions mis-regulation of the phosphorus starvation response in maize (*Zea mays* ssp. *mays* L.)"

### Supplemental Figure 1

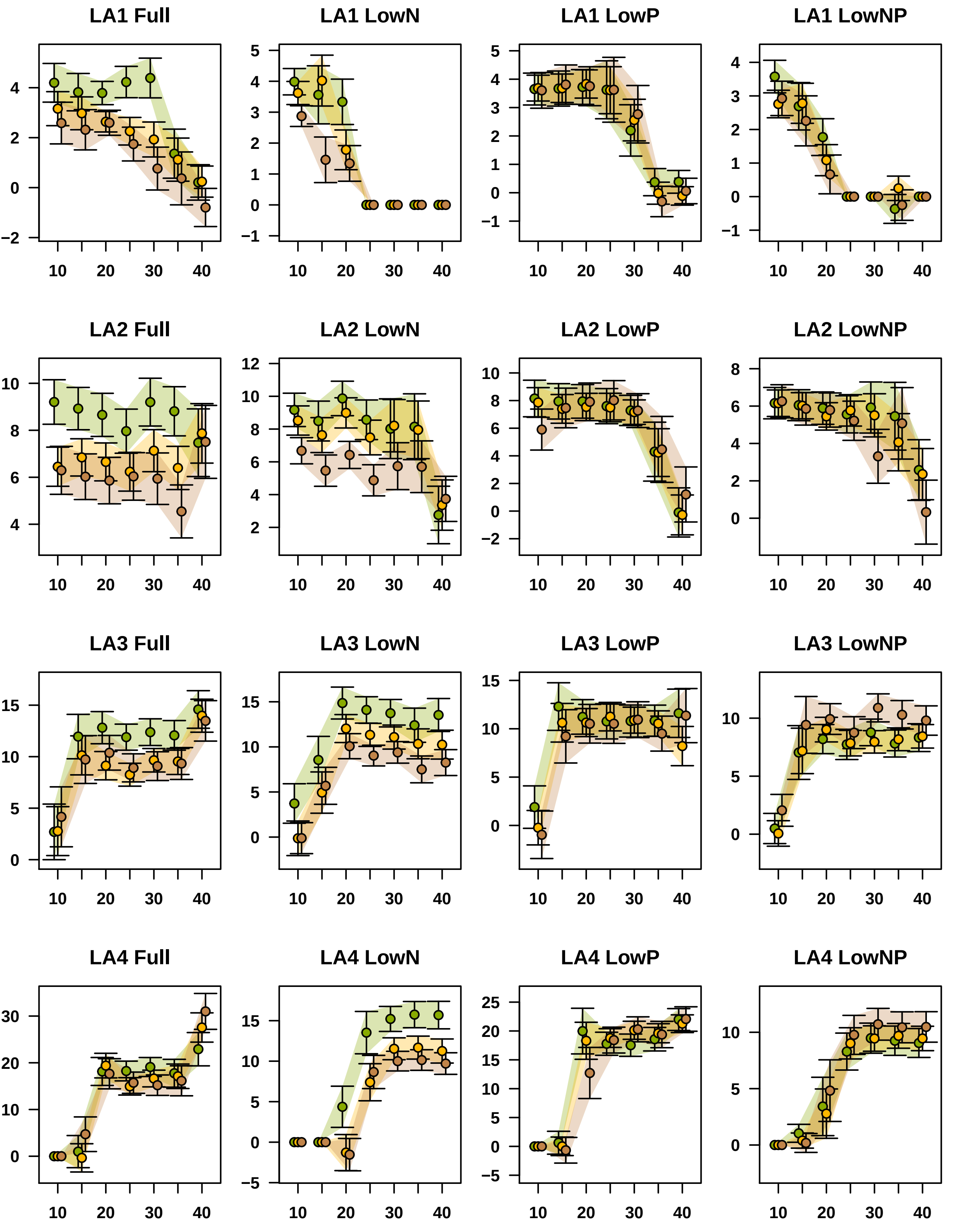

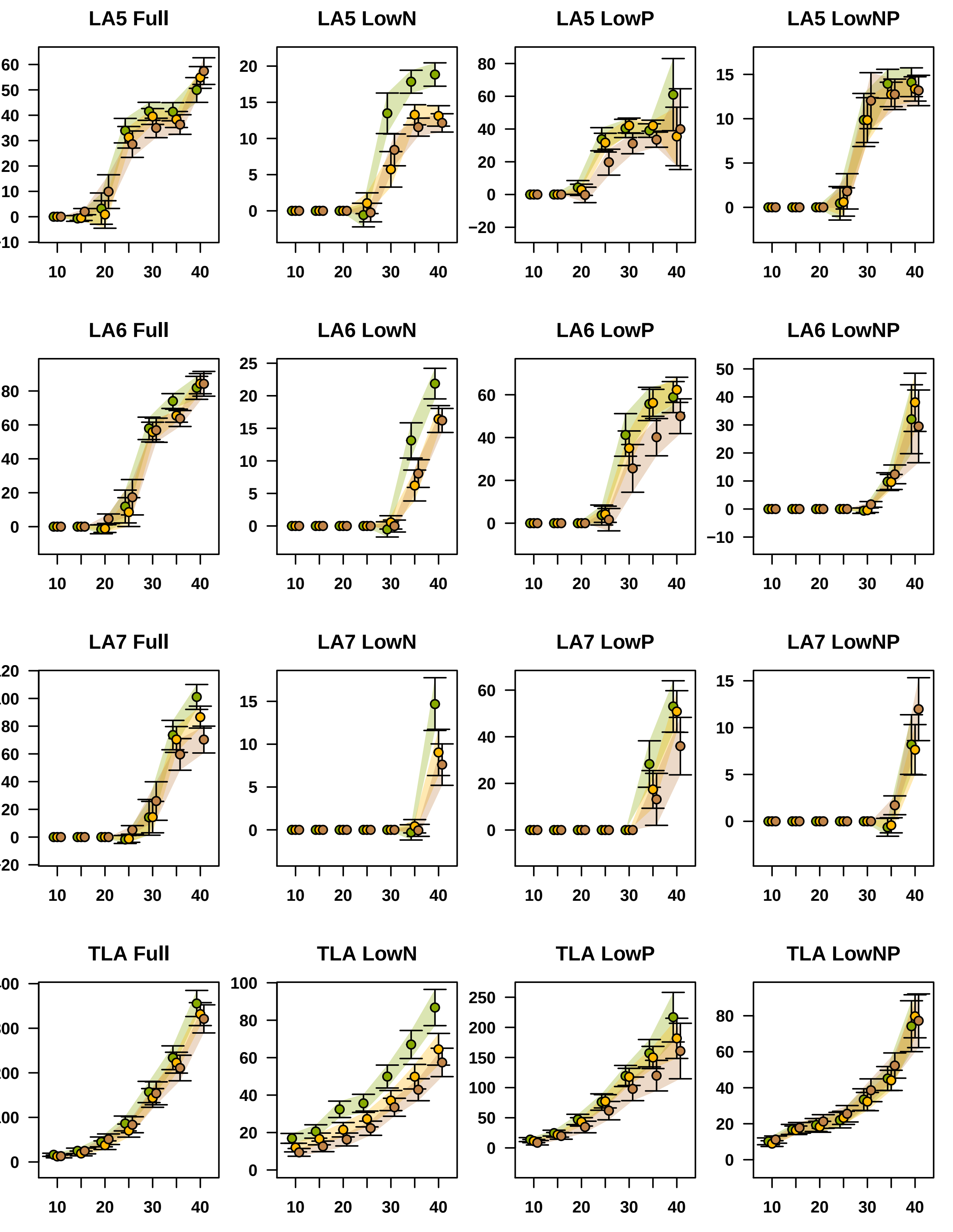

### Supplemental Figure 4

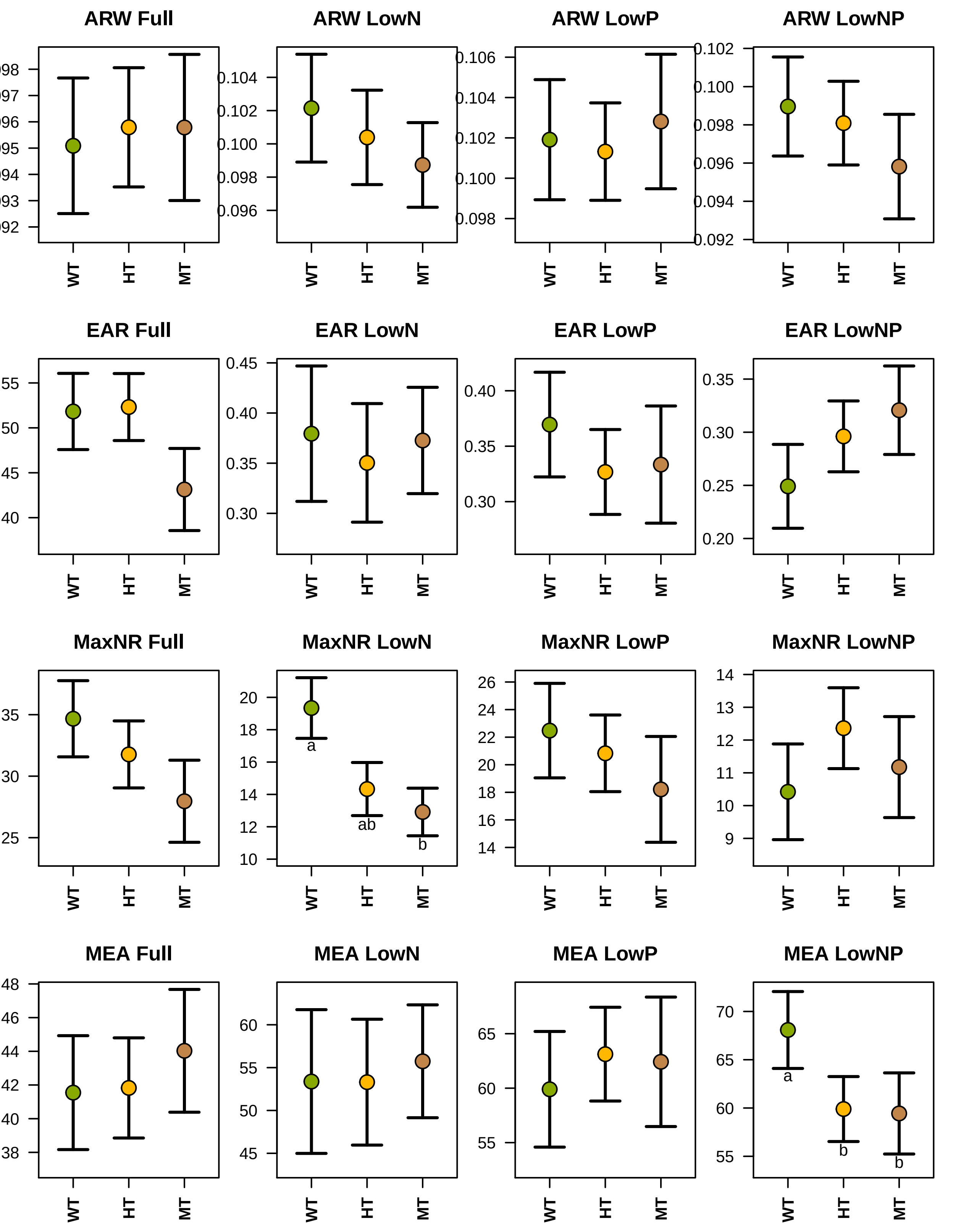

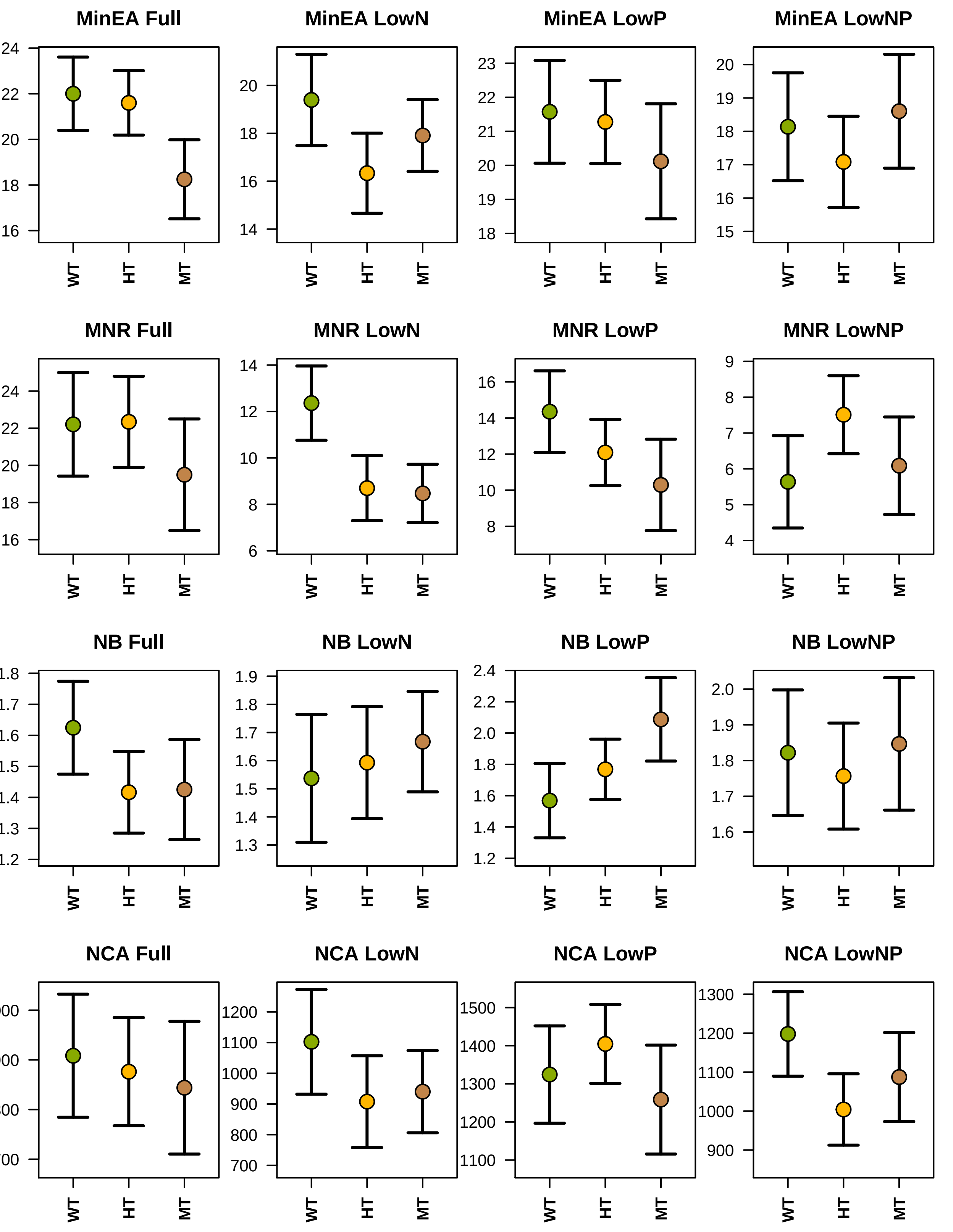

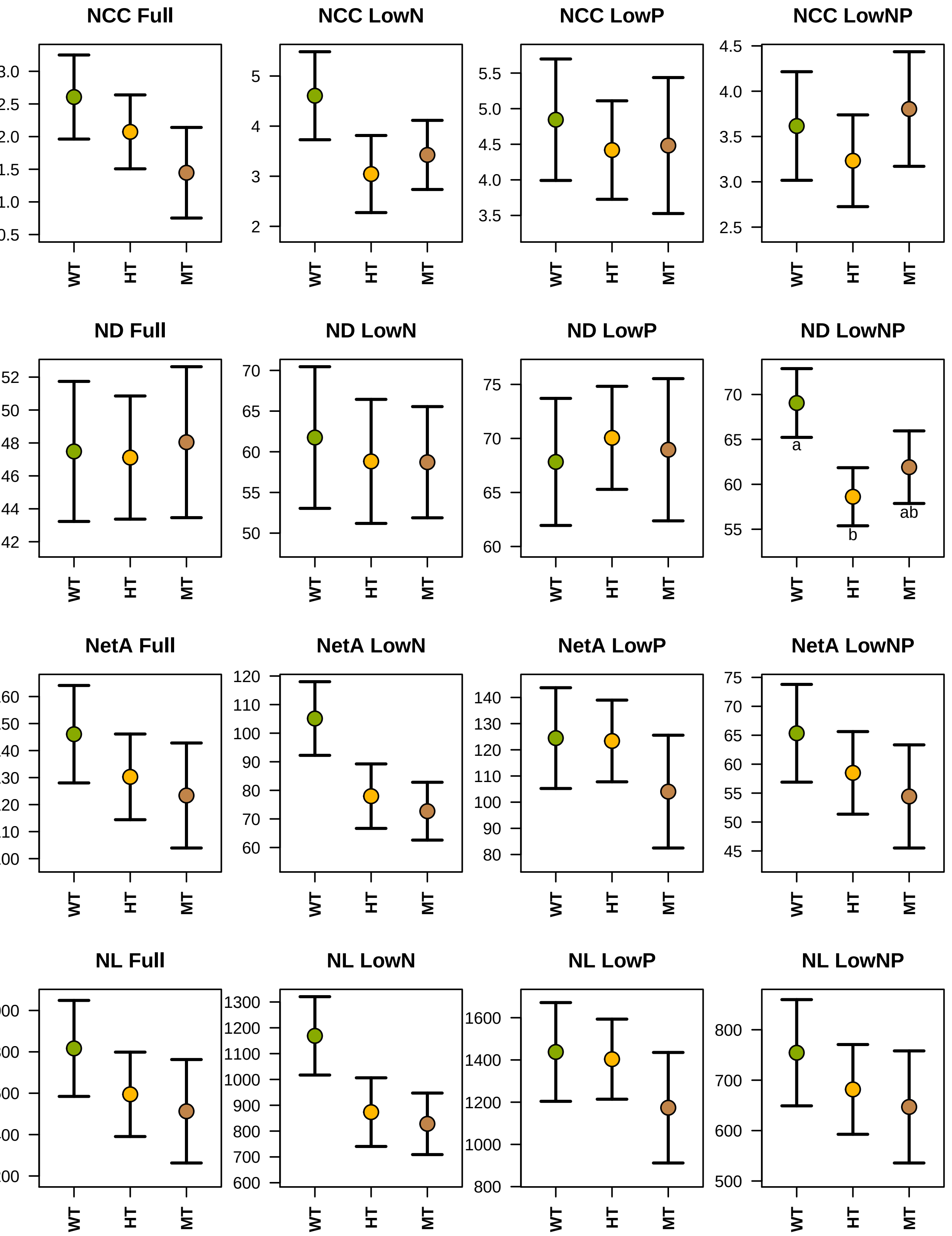

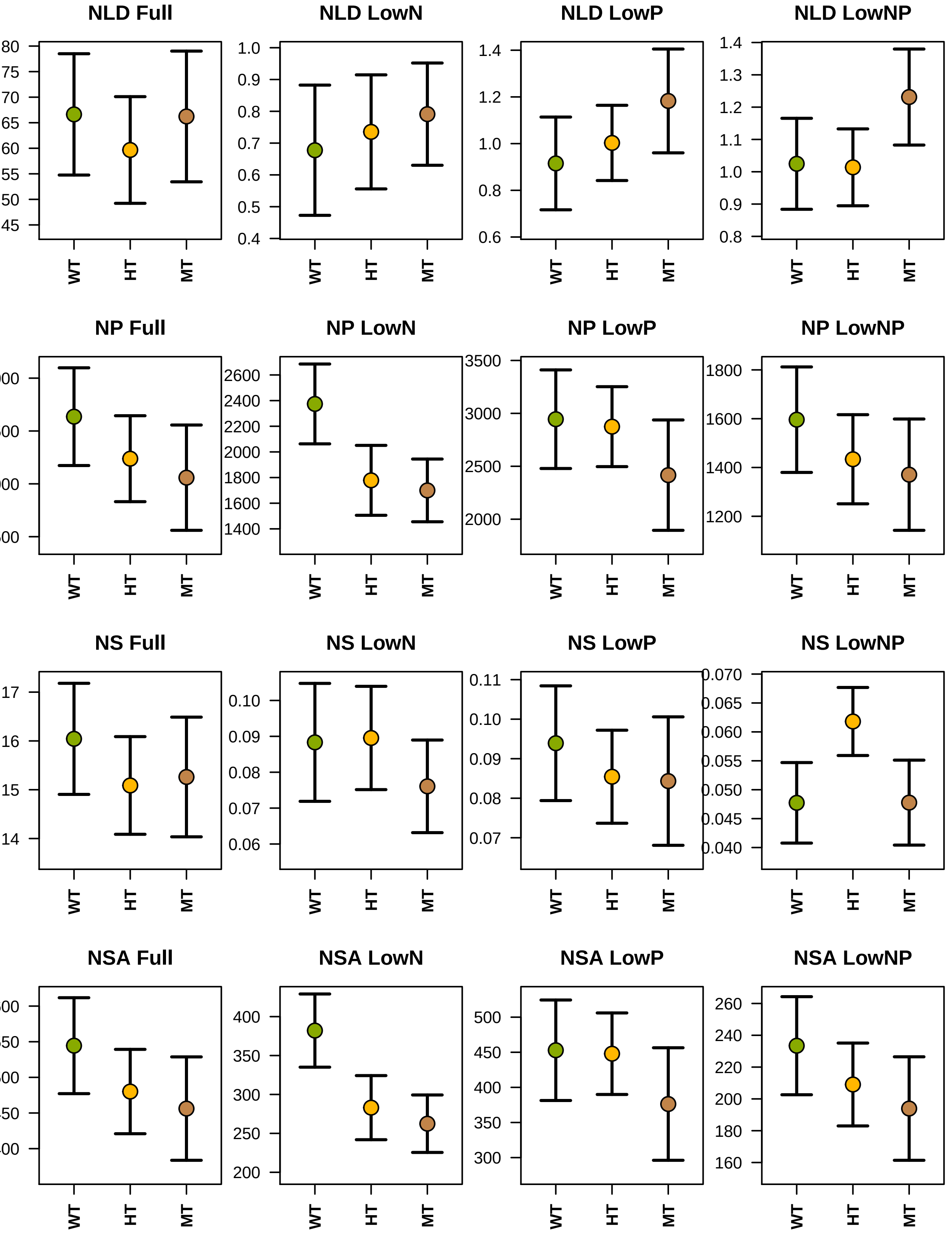

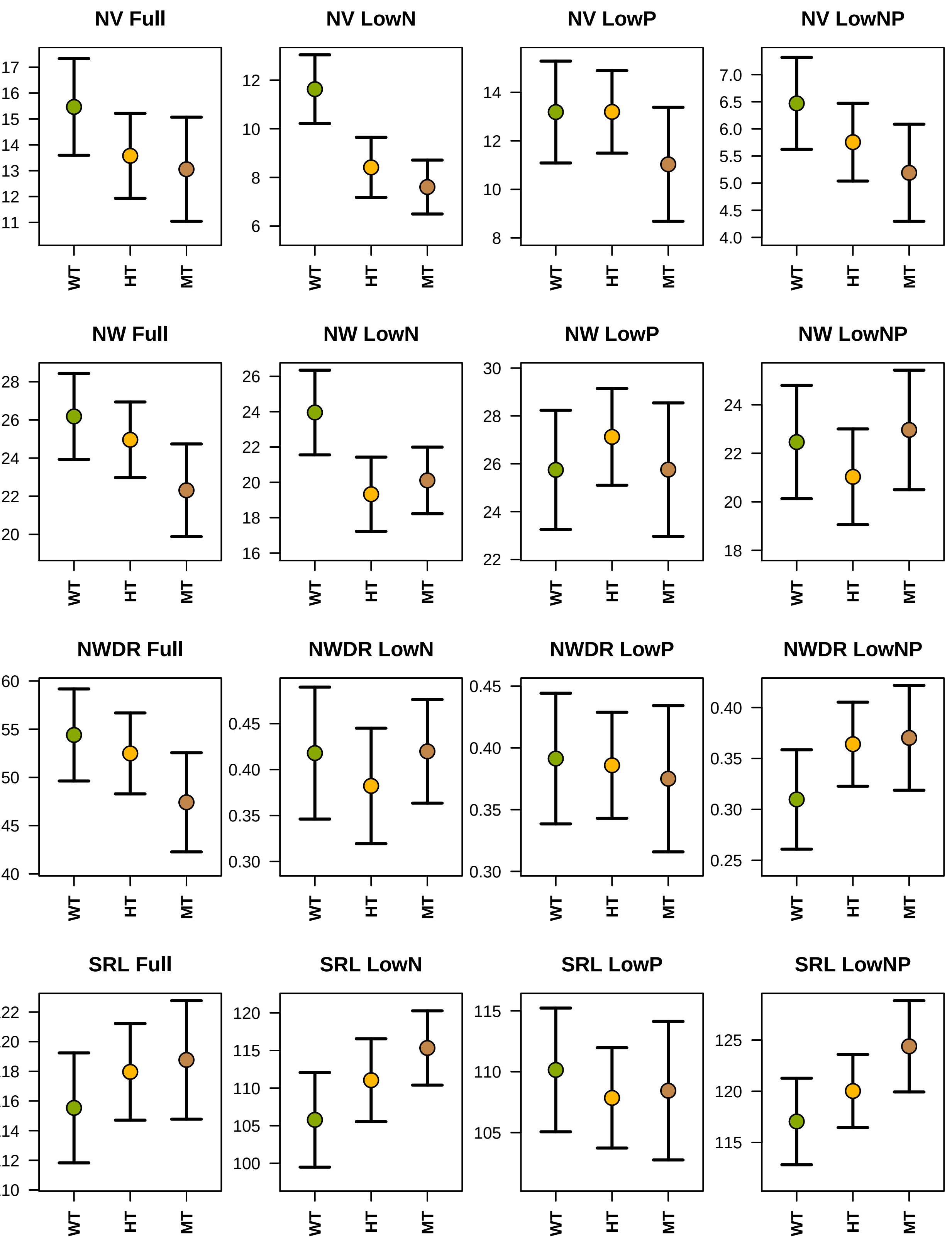
