## Supplemental Figure 2 for "The *pho1;2a’-m1.1* allele of *Phosphate1* conditions mis-regulation of the phosphorus starvation response in maize (*Zea mays* ssp. *mays* L.)"

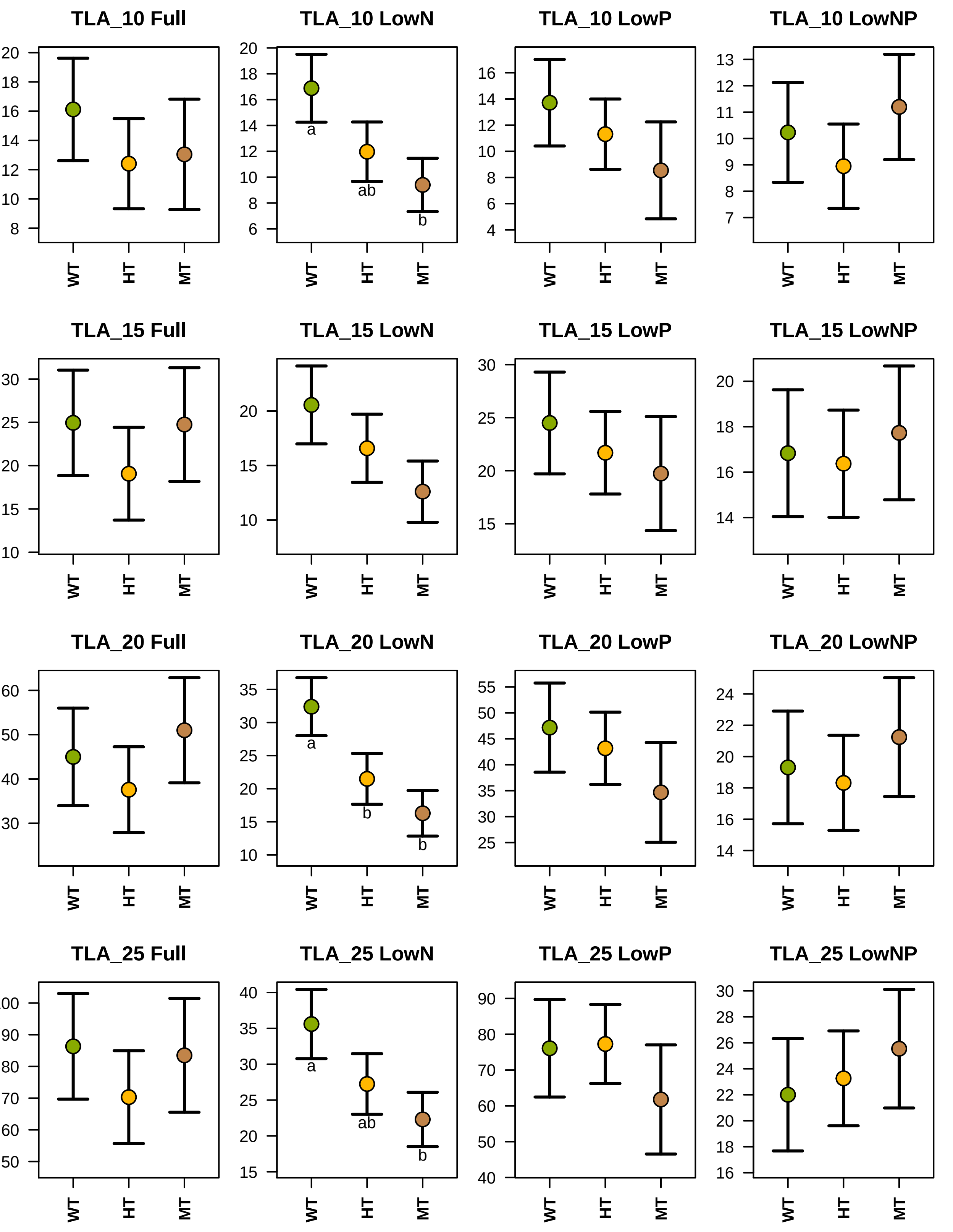
TLA_10 Full
TLA_10 LowN
TLA_10 LowP
TLA_10 LowNP
TLA_15 Full
TLA_15 LowN
TLA_15 LowP
TLA_15 LowNP
TLA_20 Full
TLA_20 LowN
TLA_20 LowP
TLA_20 LowNP
TLA_25 Full
TLA_25 LowN
TLA_25 LowP
TLA_25 LowNP
WT
HT
MT
a
ab
b

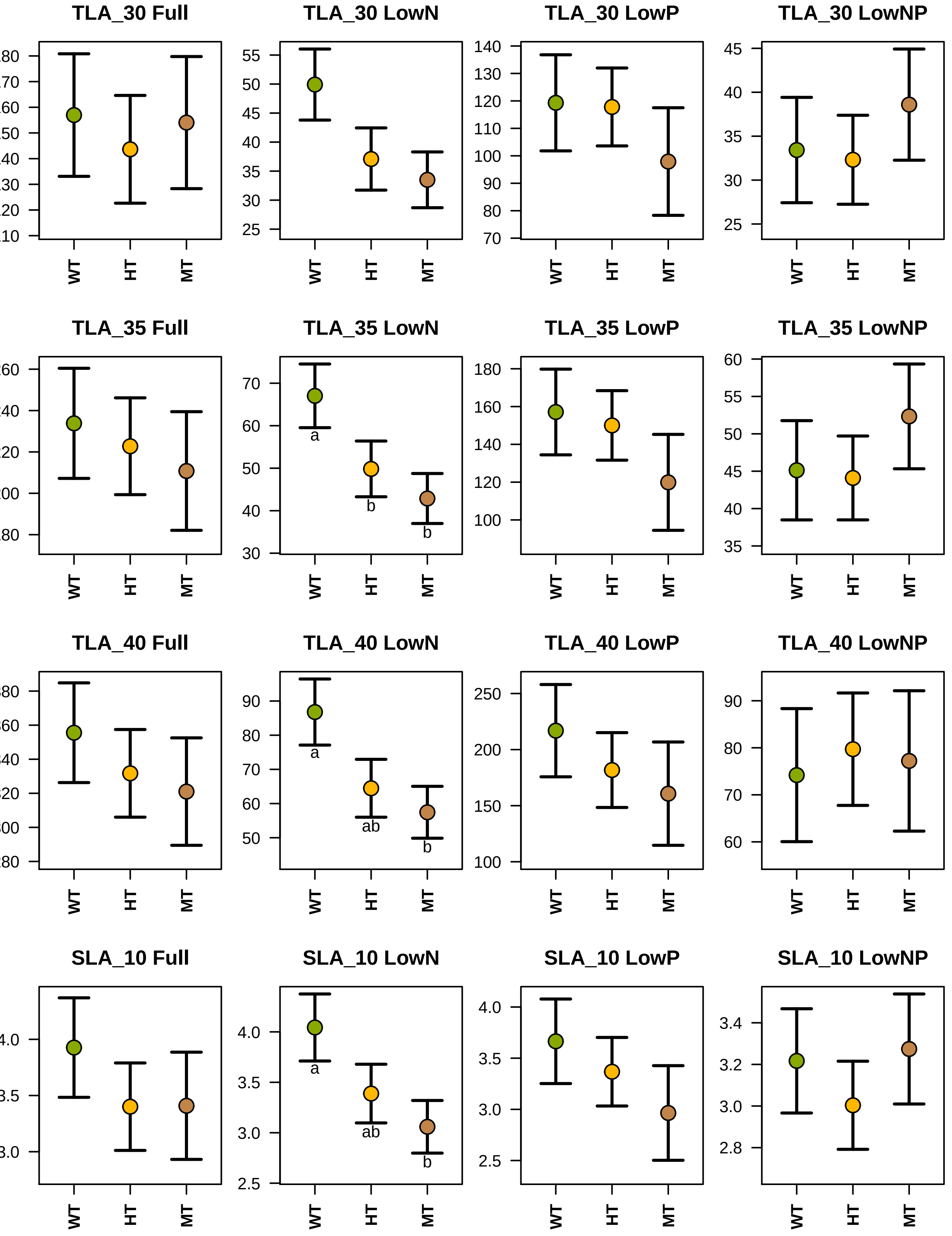

TLA_30 Full
TLA_30 LowN
TLA_30 LowP
TLA_30 LowNP
TLA_35 Full
TLA_35 LowN
TLA_35 LowP
TLA_35 LowNP
TLA_40 Full
TLA_40 LowN
TLA_40 LowP
TLA_40 LowNP
SLA_10 Full
SLA_10 LowN
SLA_10 LowP
SLA_10 LowNP
WT
HT
MT
a
b
b
a
ab
b
a
ab
b

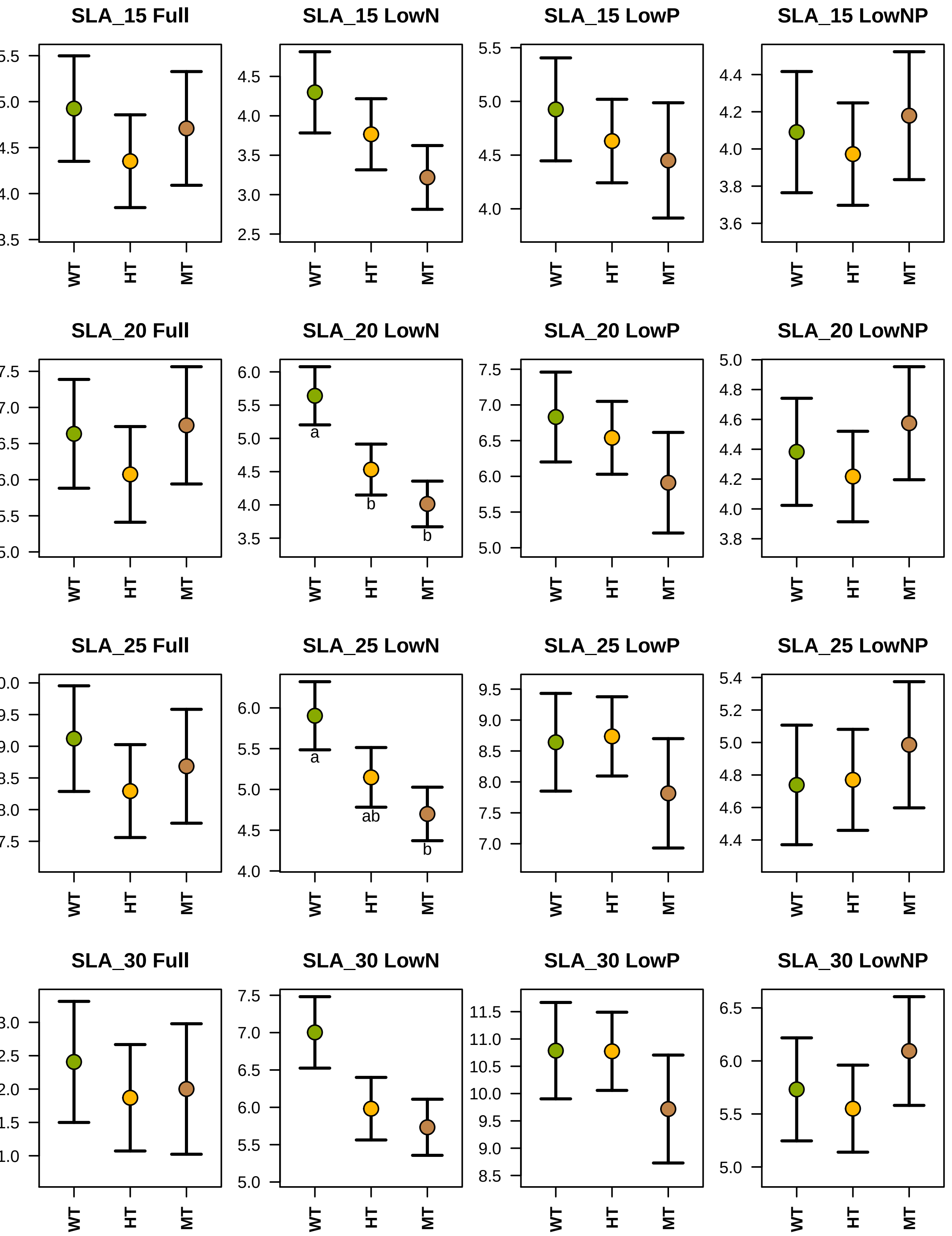

SLA_15 Full
SLA_15 LowN
4.5
4.0
3.5
3.0
2.5
SLA_15 LowP
5.5
5.0
4.5
4.0
SLA_15 LowNP
4.4
4.2
4.0
3.8
3.6
SLA_20 Full
SLA_20 LowN
6.0
5.5
5.0
4.5
4.0
3.5
a
b
b
SLA_20 LowP
7.5
7.0
6.5
6.0
5.5
5.0
SLA_20 LowNP
5.0
4.8
4.6
4.4
4.2
4.0
3.8
SLA_25 Full
SLA_25 LowN
6.0
5.5
5.0
4.5
4.0
a
ab
b
SLA_25 LowP
9.5
9.0
8.5
8.0
7.5
7.0
SLA_25 LowNP
5.4
5.2
5.0
4.8
4.6
4.4
SLA_30 Full
SLA_30 LowN
7.5
7.0
6.5
6.0
5.5
5.0
SLA_30 LowP
11.5
11.0
10.5
10.0
9.5
9.0
8.5
SLA_30 LowNP
6.5
6.0
5.5
5.0
WT
HT
MT

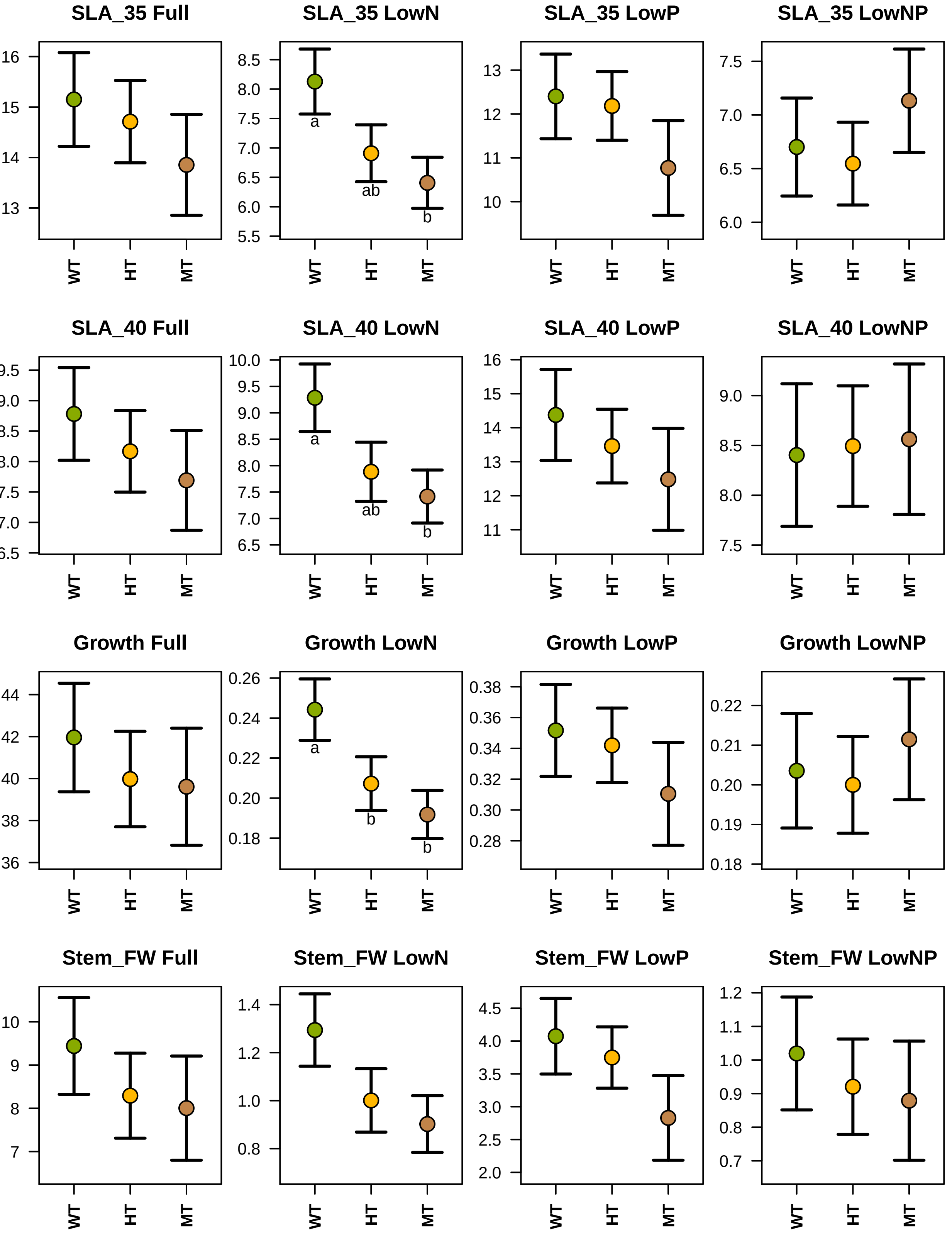
SLA_35 Full
SLA_35 LowN
SLA_35 LowP
SLA_35 LowNP
SLA_40 Full
SLA_40 LowN
SLA_40 LowP
SLA_40 LowNP
Growth Full
Growth LowN
Growth LowP
Growth LowNP
Stem_FW Full
Stem_FW LowN
Stem_FW LowP
Stem_FW LowNP
WT
HT
MT
a
ab
b

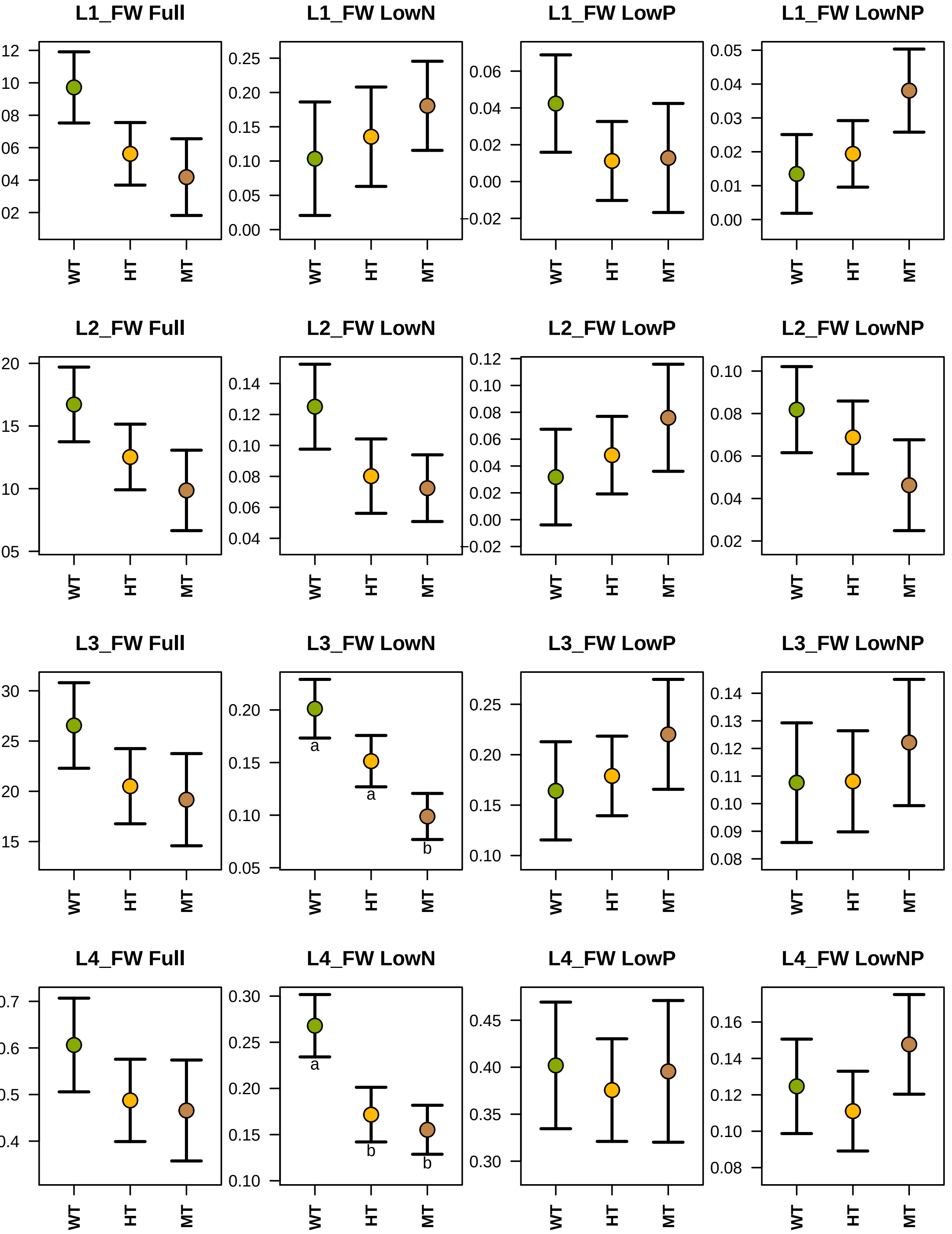

L1_FW Full
L1_FW LowN
L1_FW LowP
L1_FW LowNP
L2_FW Full
L2_FW LowN
L2_FW LowP
L2_FW LowNP
L3_FW Full
L3_FW LowN
L3_FW LowP
L3_FW LowNP
L4_FW Full
L4_FW LowN
L4_FW LowP
L4_FW LowNP
WT
HT
MT
a
b

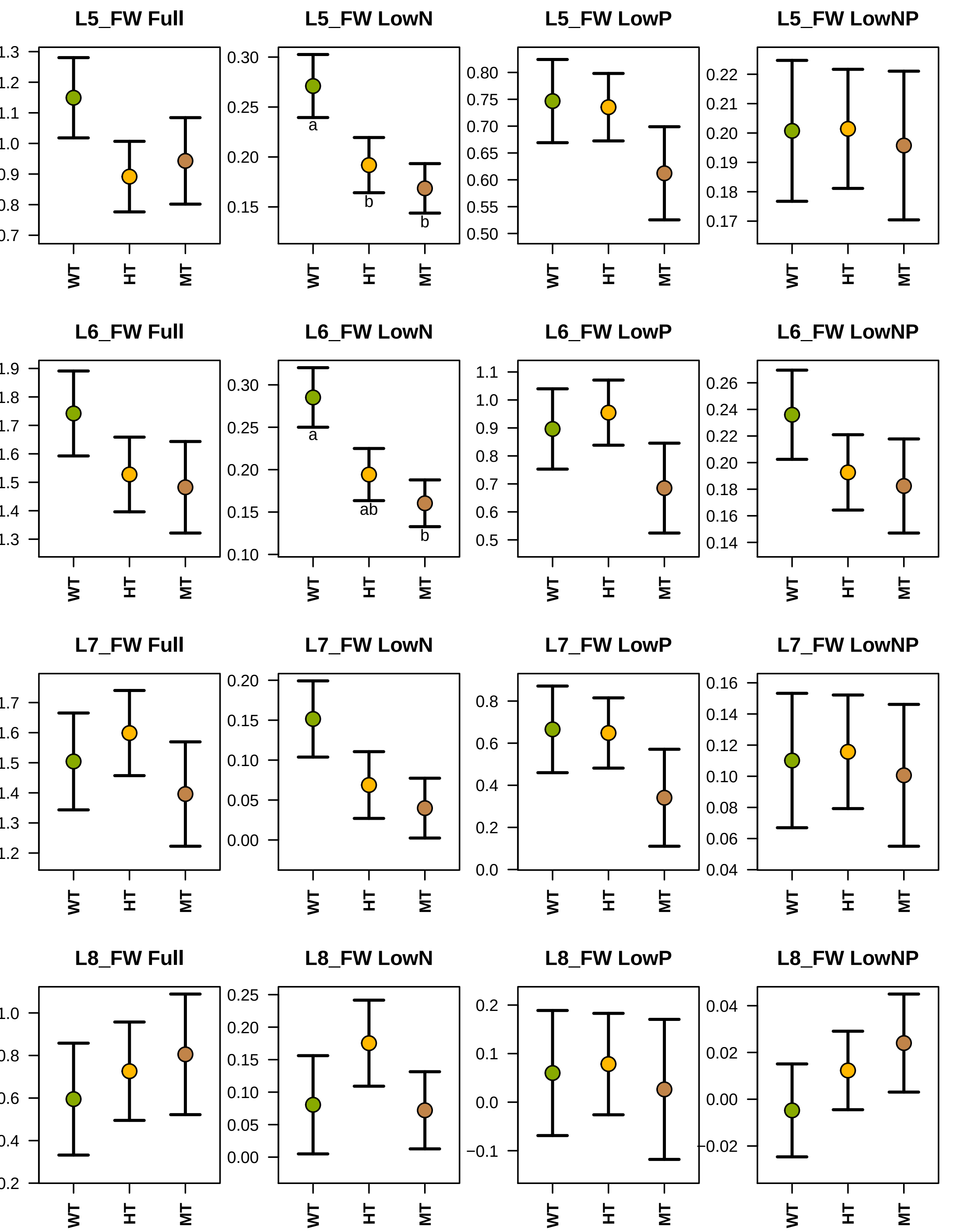

L5_FW Full
L5_FW LowN
L5_FW LowP
L5_FW LowNP
L6_FW Full
L6_FW LowN
L6_FW LowP
L6_FW LowNP
L7_FW Full
L7_FW LowN
L7_FW LowP
L7_FW LowNP
L8_FW Full
L8_FW LowN
L8_FW LowP
L8_FW LowNP
a
b
b
a
ab
b
WT
HT
MT

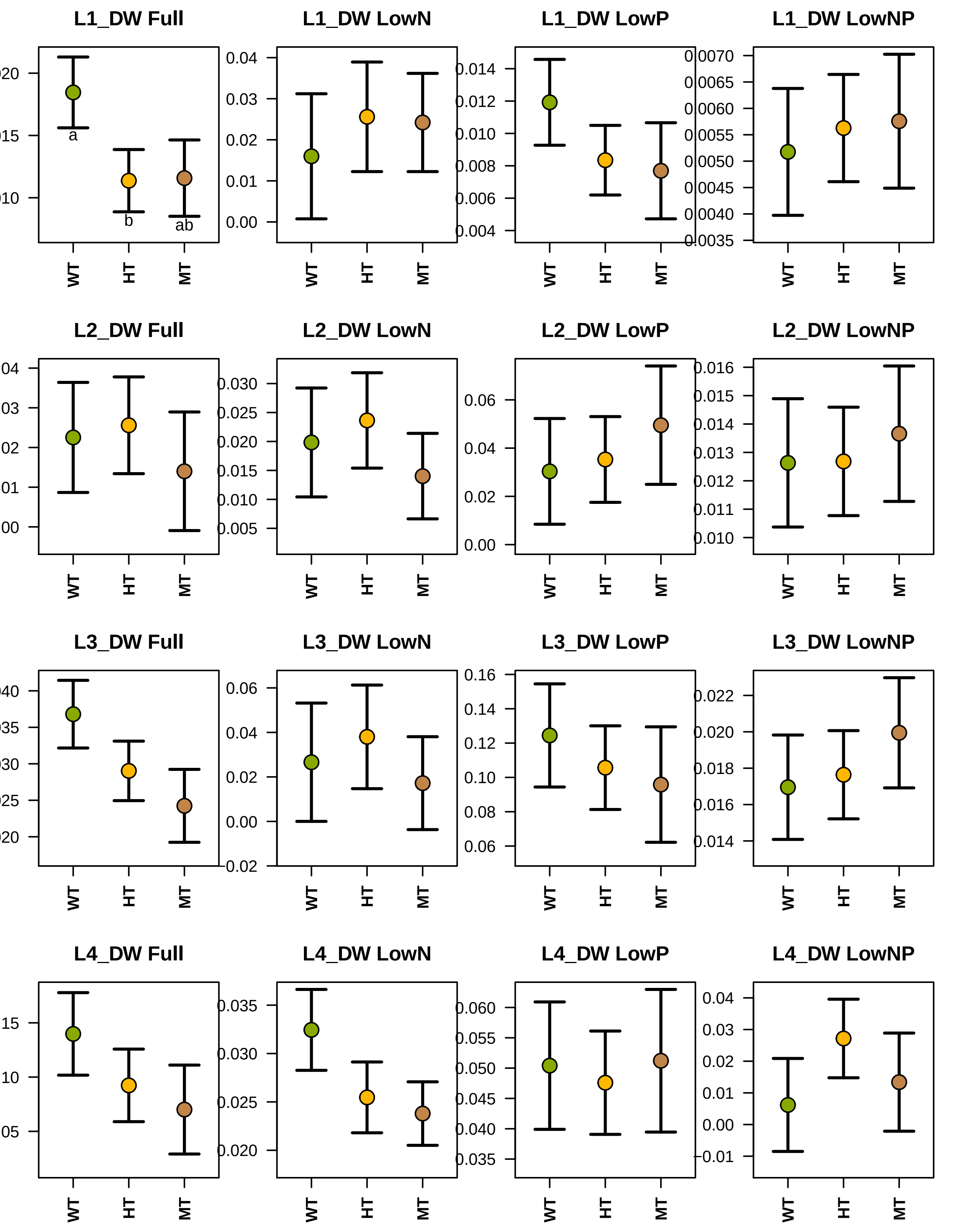
L1_DW Full
a
b
ab
WT HT MT
L1_DW LowN
0.00 0.01 0.02 0.03 0.04
WT HT MT
L1_DW LowP
0.004 0.006 0.008 0.010 0.012 0.014
WT HT MT
L1_DW LowNP
0.0040 0.0045 0.0050 0.0055 0.0060 0.0065 0.0070
WT HT MT
L2_DW Full
WT HT MT
L2_DW LowN
0.005 0.010 0.015 0.020 0.025 0.030
WT HT MT
L2_DW LowP
0.00 0.02 0.04 0.06
WT HT MT
L2_DW LowNP
0.010 0.011 0.012 0.013 0.014 0.015 0.016
WT HT MT
L3_DW Full
WT HT MT
L3_DW LowN
-0.02 0.00 0.02 0.04 0.06
WT HT MT
L3_DW LowP
0.06 0.08 0.10 0.12 0.14 0.16
WT HT MT
L3_DW LowNP
0.014 0.016 0.018 0.020 0.022
WT HT MT
L4_DW Full
WT HT MT
L4_DW LowN
0.020 0.025 0.030 0.035
WT HT MT
L4_DW LowP
0.035 0.040 0.045 0.050 0.055 0.060
WT HT MT
L4_DW LowNP
-0.01 0.00 0.01 0.02 0.03 0.04
WT HT MT

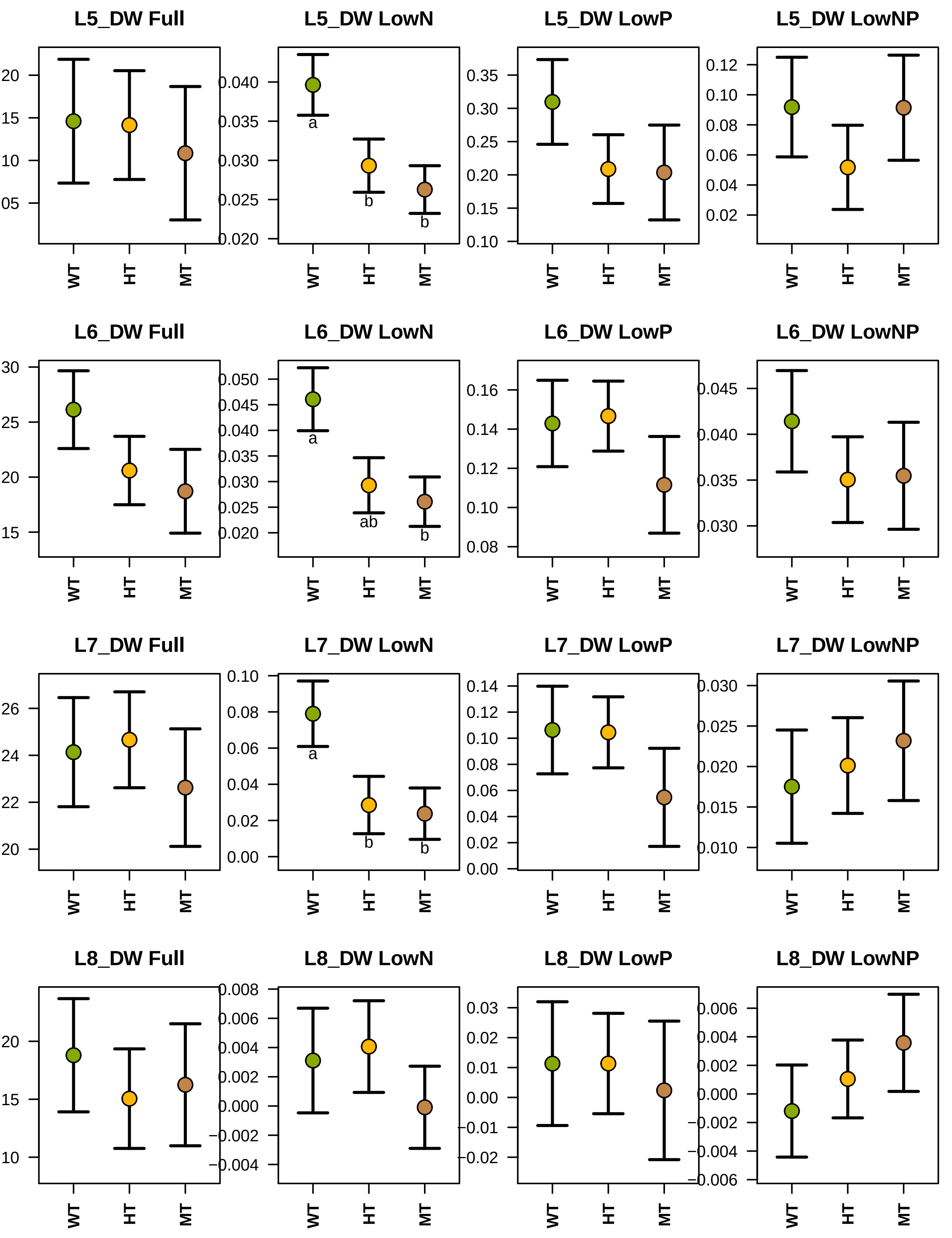

L5_DW Full
20
15
10
05
WT
HT
MT
L5_DW LowN
0.040
0.035
0.030
0.025
0.020
a
b
b
WT
HT
MT
L5_DW LowP
0.35
0.30
0.25
0.20
0.15
0.10
WT
HT
MT
L5_DW LowNP
0.12
0.10
0.08
0.06
0.04
0.02
WT
HT
MT
L6_DW Full
30
25
20
15
WT
HT
MT
L6_DW LowN
0.050
0.045
0.040
0.035
0.030
0.025
0.020
a
ab
b
WT
HT
MT
L6_DW LowP
0.16
0.14
0.12
0.10
0.08
WT
HT
MT
L6_DW LowNP
0.045
0.040
0.035
0.030
WT
HT
MT
L7_DW Full
26
24
22
20
WT
HT
MT
L7_DW LowN
0.10
0.08
0.06
0.04
0.02
0.00
a
b
b
WT
HT
MT
L7_DW LowP
0.14
0.12
0.10
0.08
0.06
0.04
0.02
0.00
WT
HT
MT
L7_DW LowNP
0.030
0.025
0.020
0.015
0.010
WT
HT
MT
L8_DW Full
20
15
10
WT
HT
MT
L8_DW LowN
0.008
0.006
0.004
0.002
0.000
−0.002
−0.004
WT
HT
MT
L8_DW LowP
0.03
0.02
0.01
0.00
−0.01
−0.02
WT
HT
MT
L8_DW LowNP
0.006
0.004
0.002
0.000
−0.002
−0.004
−0.006
WT
HT
MT

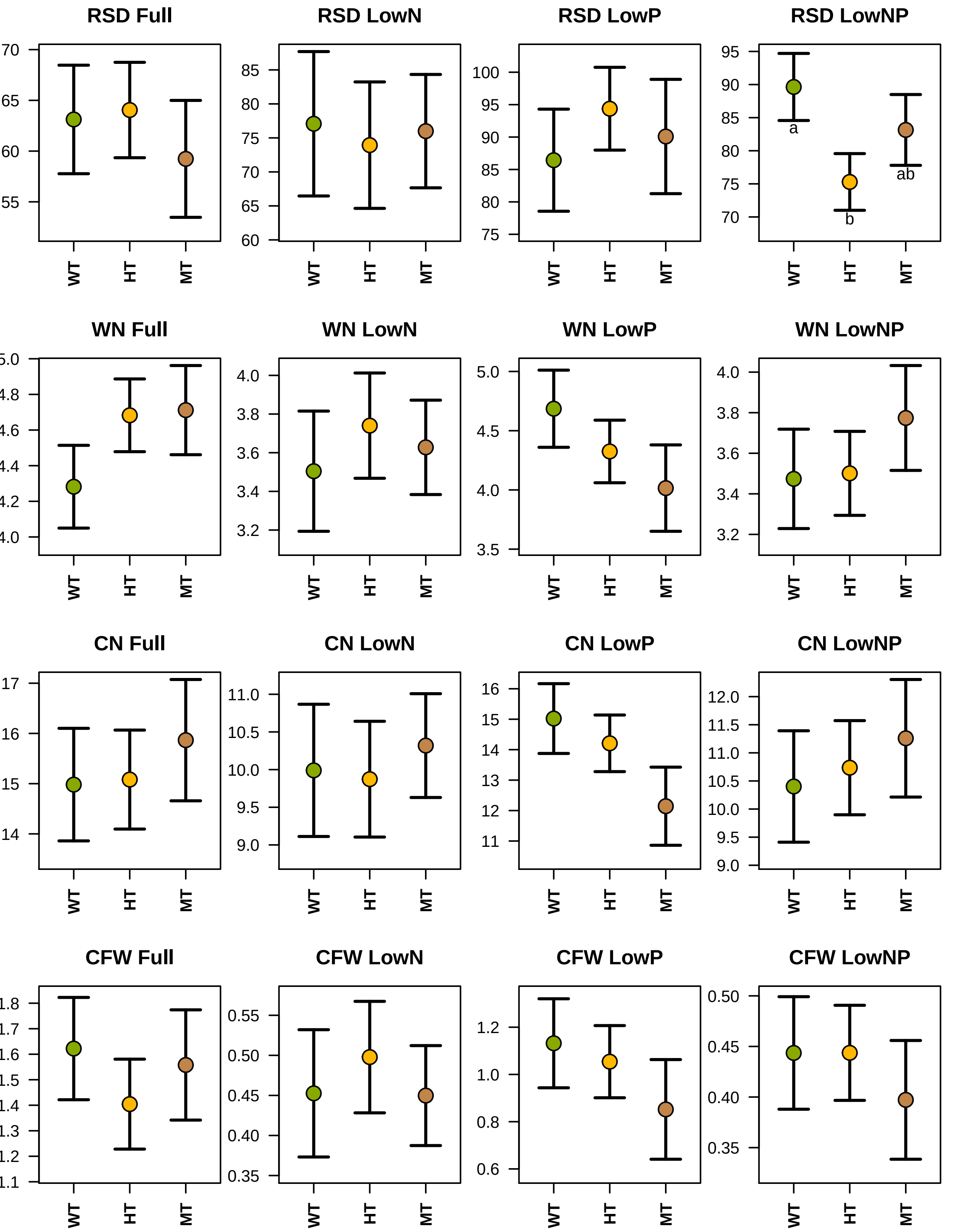

RSD Full
RSD LowN
RSD LowP
RSD LowNP
a
b
ab
WN Full
WN LowN
WN LowP
WN LowNP
CN Full
CN LowN
CN LowP
CN LowNP
CFW Full
CFW LowN
CFW LowP
CFW LowNP
WT
HT
MT

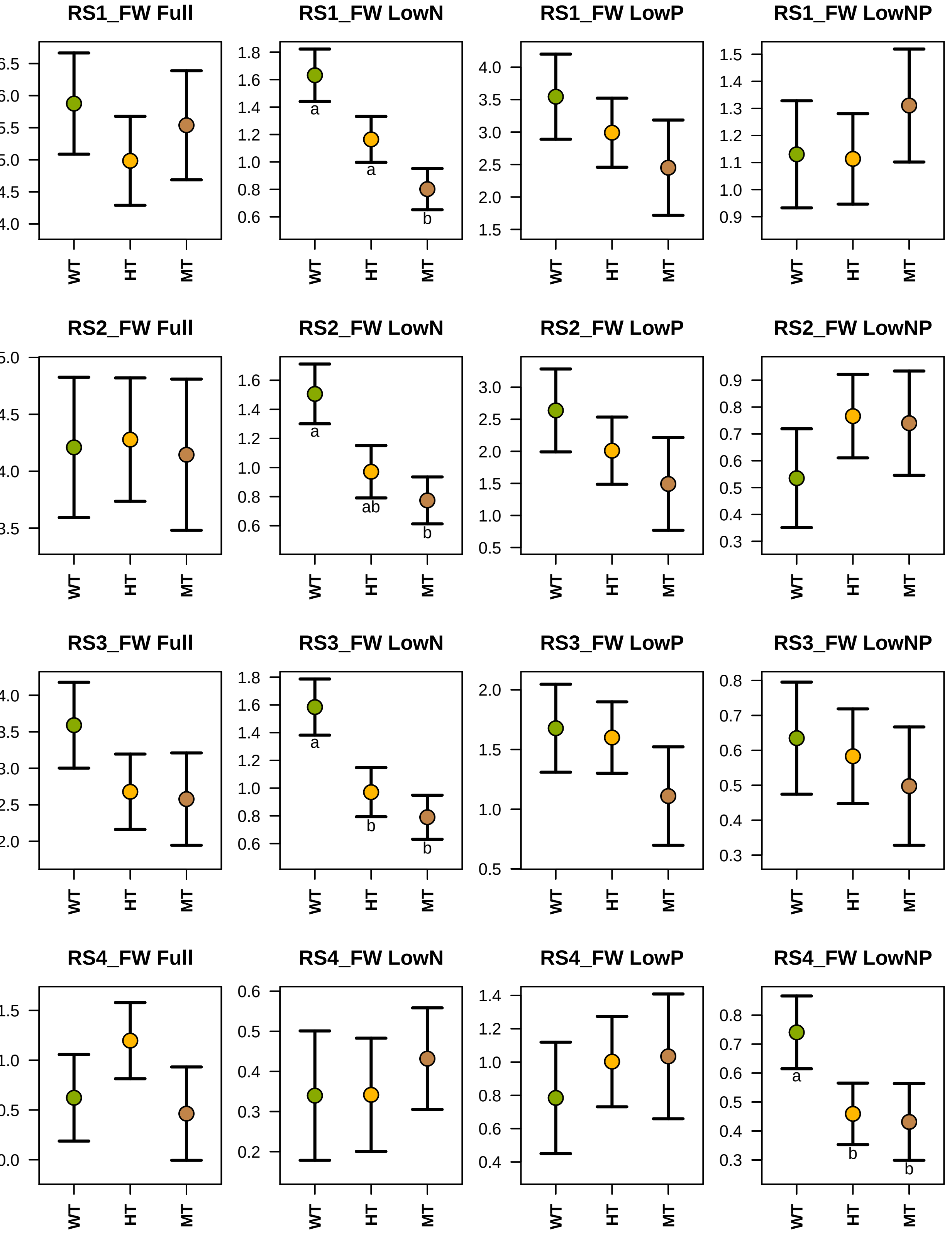
RS1_FW Full
RS1_FW LowN
a
a
b
RS1_FW LowP
RS1_FW LowNP
RS2_FW Full
RS2_FW LowN
a
ab
b
RS2_FW LowP
RS2_FW LowNP
RS3_FW Full
RS3_FW LowN
a
b
b
RS3_FW LowP
RS3_FW LowNP
RS4_FW Full
RS4_FW LowN
RS4_FW LowP
RS4_FW LowNP
a
b
b
WT
HT
MT

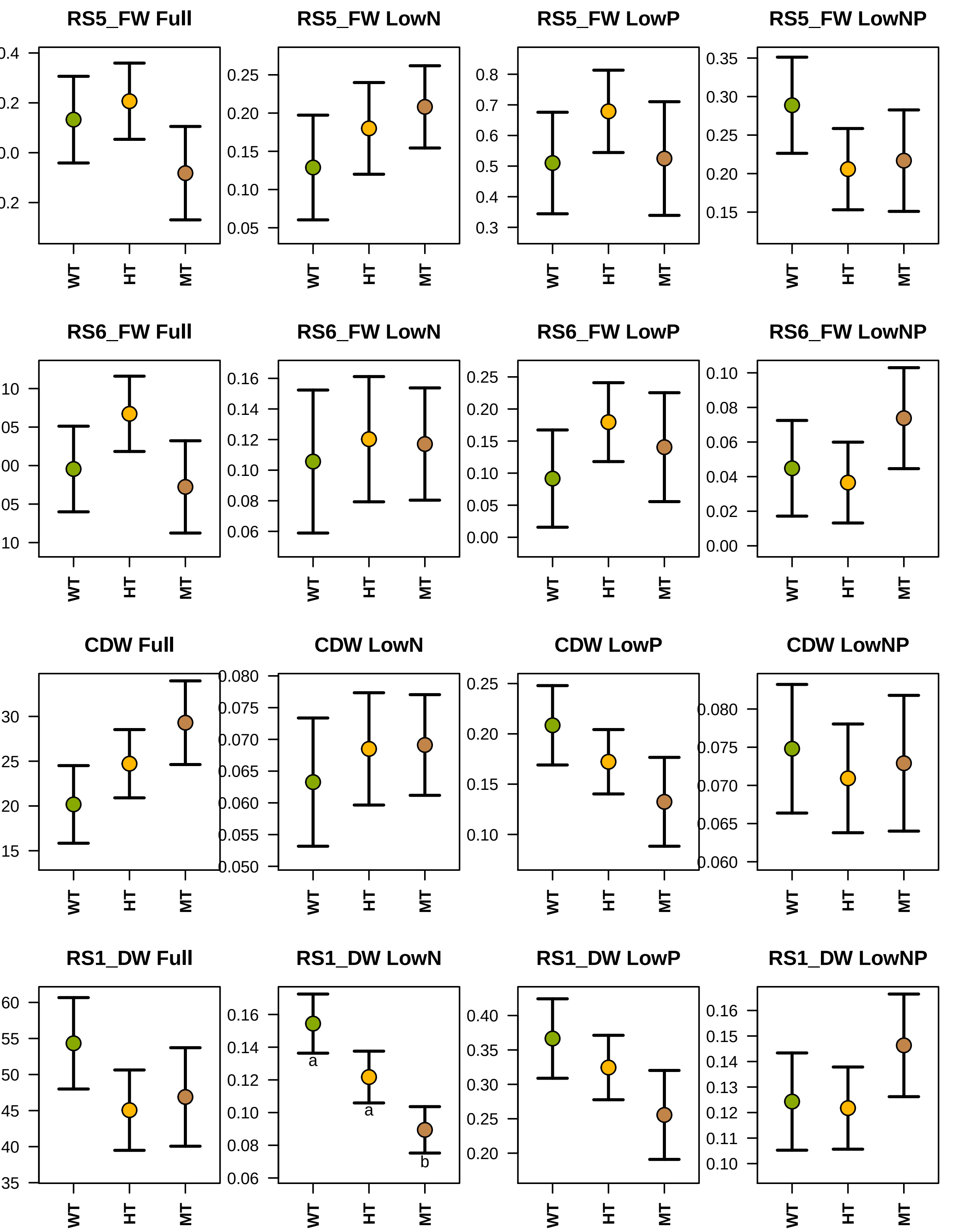
RS5_FW Full
RS5_FW LowN
RS5_FW LowP
RS5_FW LowNP
RS6_FW Full
RS6_FW LowN
RS6_FW LowP
RS6_FW LowNP
CDW Full
CDW LowN
CDW LowP
CDW LowNP
RS1_DW Full
RS1_DW LowN
RS1_DW LowP
RS1_DW LowNP
WT
HT
MT
a
a
b

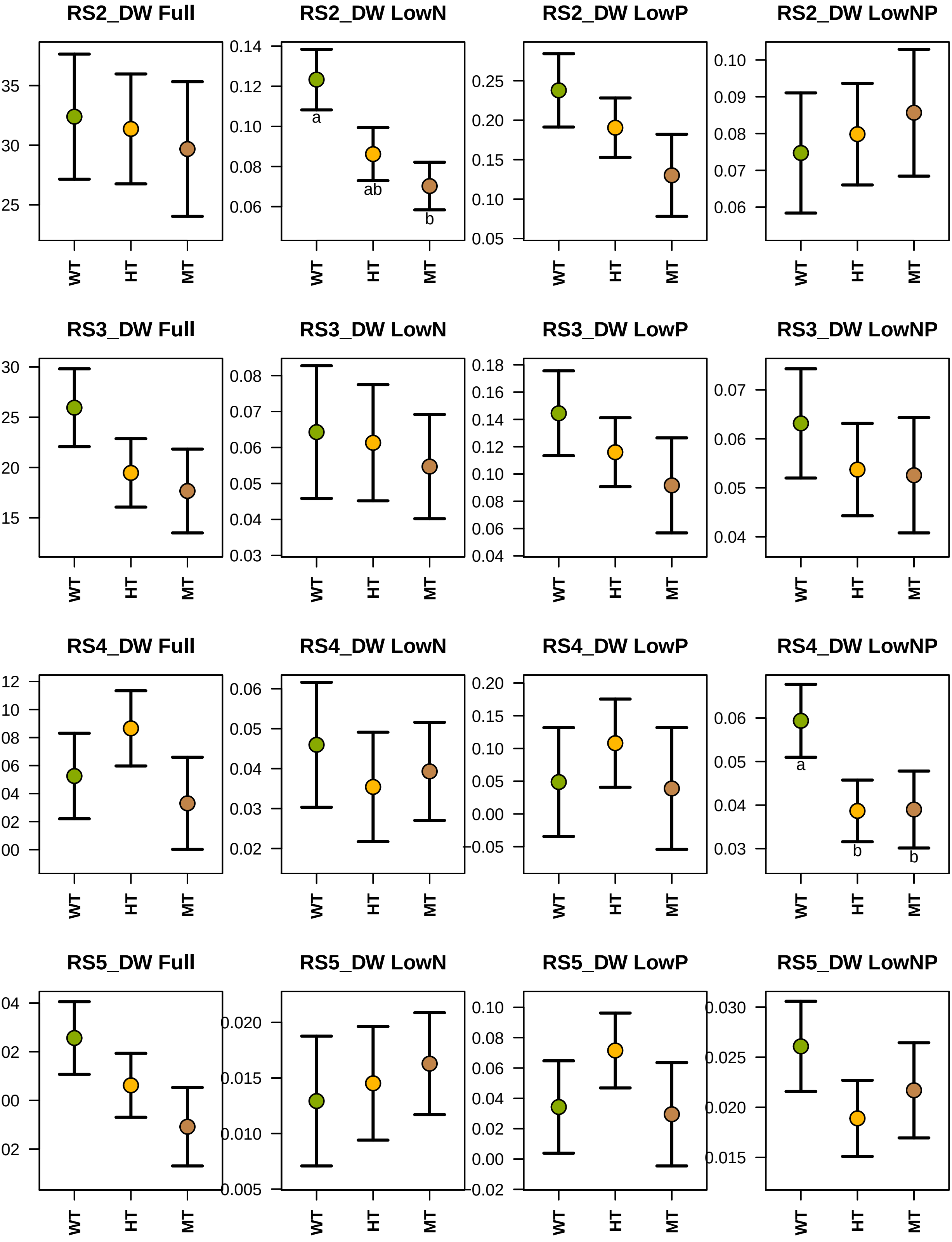
RS2_DW Full
35
30
25
WT
HT
MT
RS2_DW LowN
0.14
0.12
0.10
0.08
0.06
a
ab
b
WT
HT
MT
RS2_DW LowP
0.25
0.20
0.15
0.10
0.05
WT
HT
MT
RS2_DW LowNP
0.10
0.09
0.08
0.07
0.06
WT
HT
MT
RS3_DW Full
30
25
20
15
WT
HT
MT
RS3_DW LowN
0.08
0.07
0.06
0.05
0.04
0.03
WT
HT
MT
RS3_DW LowP
0.18
0.16
0.14
0.12
0.10
0.08
0.06
0.04
WT
HT
MT
RS3_DW LowNP
0.07
0.06
0.05
0.04
WT
HT
MT
RS4_DW Full
12
10
08
06
04
02
00
WT
HT
MT
RS4_DW LowN
0.06
0.05
0.04
0.03
0.02
WT
HT
MT
RS4_DW LowP
0.20
0.15
0.10
0.05
0.00
-0.05
WT
HT
MT
RS4_DW LowNP
0.06
0.05
0.04
0.03
a
b
b
WT
HT
MT
RS5_DW Full
04
02
00
02
WT
HT
MT
RS5_DW LowN
0.020
0.015
0.010
0.005
WT
HT
MT
RS5_DW LowP
0.10
0.08
0.06
0.04
0.02
0.00
-0.02
WT
HT
MT
RS5_DW LowNP
0.030
0.025
0.020
0.015
WT
HT
MT

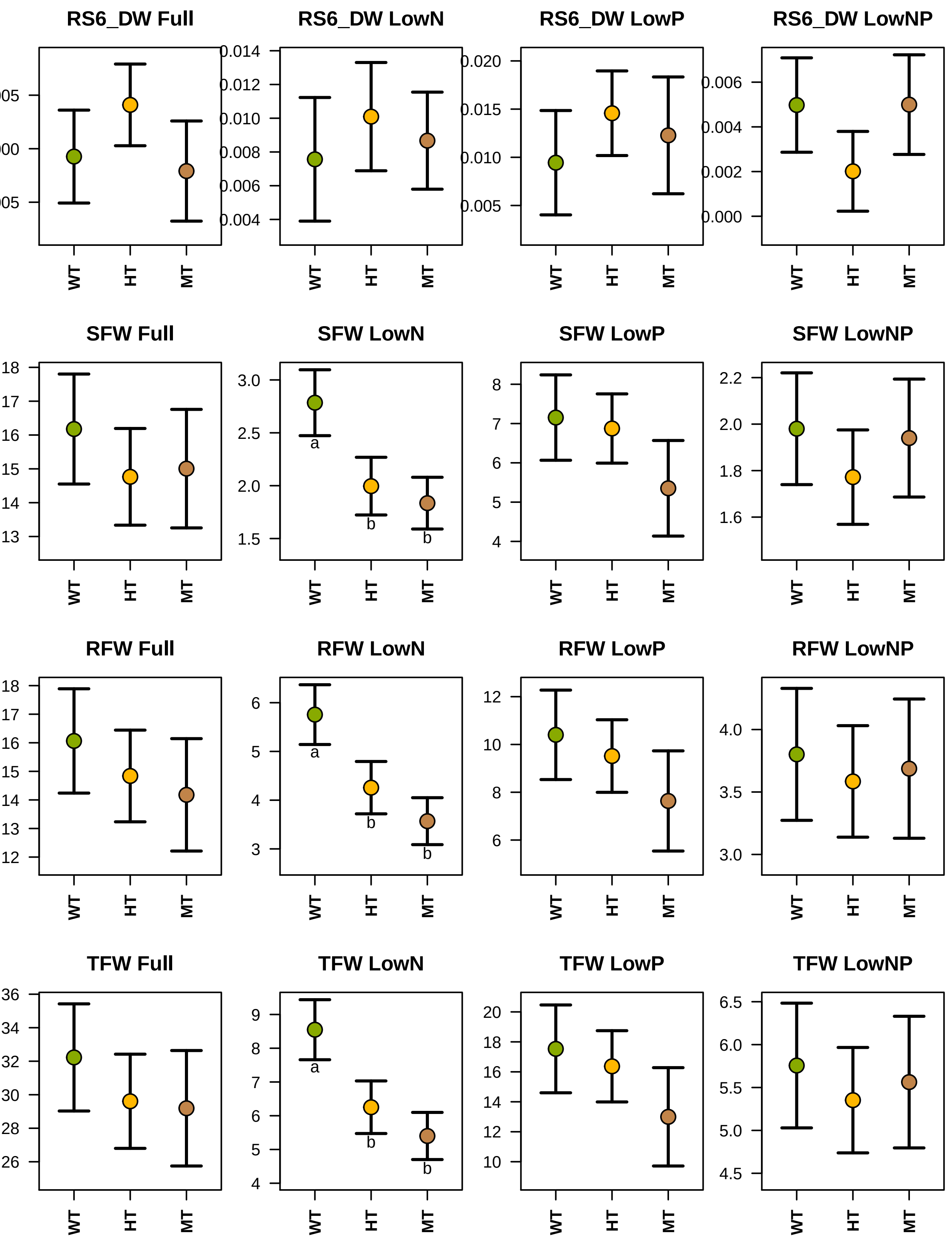

RS6_DW Full
RS6_DW LowN
RS6_DW LowP
RS6_DW LowNP
SFW Full
SFW LowN
SFW LowP
SFW LowNP
RFW Full
RFW LowN
RFW LowP
RFW LowNP
TFW Full
TFW LowN
TFW LowP
TFW LowNP
WT
HT
MT
a
b
b

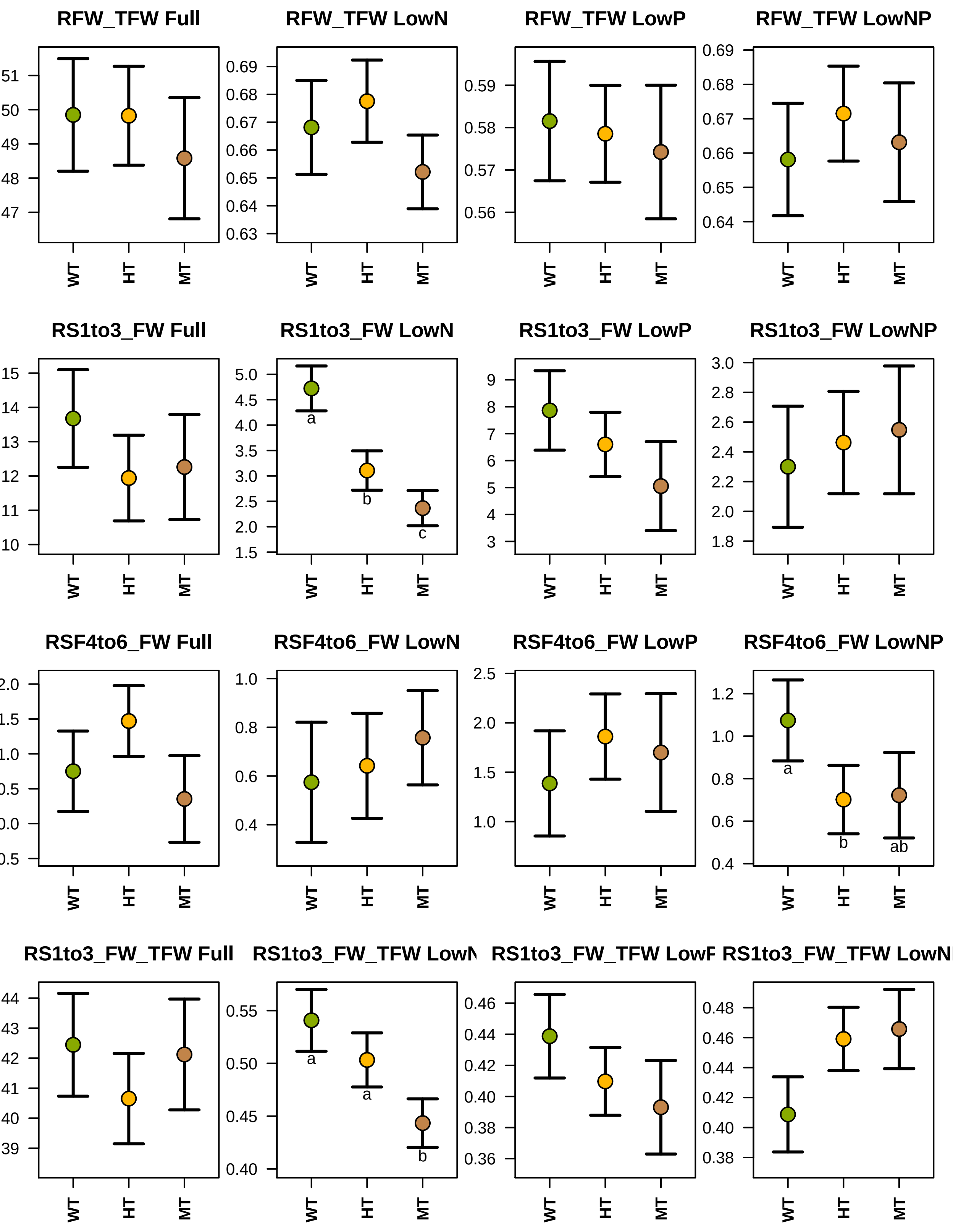

RFW_TFW Full
RFW_TFW LowN
RFW_TFW LowP
RFW_TFW LowNP
RS1to3_FW Full
RS1to3_FW LowN
RS1to3_FW LowP
RS1to3_FW LowNP
RSF4to6_FW Full
RSF4to6_FW LowN
RSF4to6_FW LowP
RSF4to6_FW LowNP
RS1to3_FW_TFW Full
RS1to3_FW_TFW LowN
RS1to3_FW_TFW LowP
RS1to3_FW_TFW LowNP
WT
HT
MT
a
b
c
ab

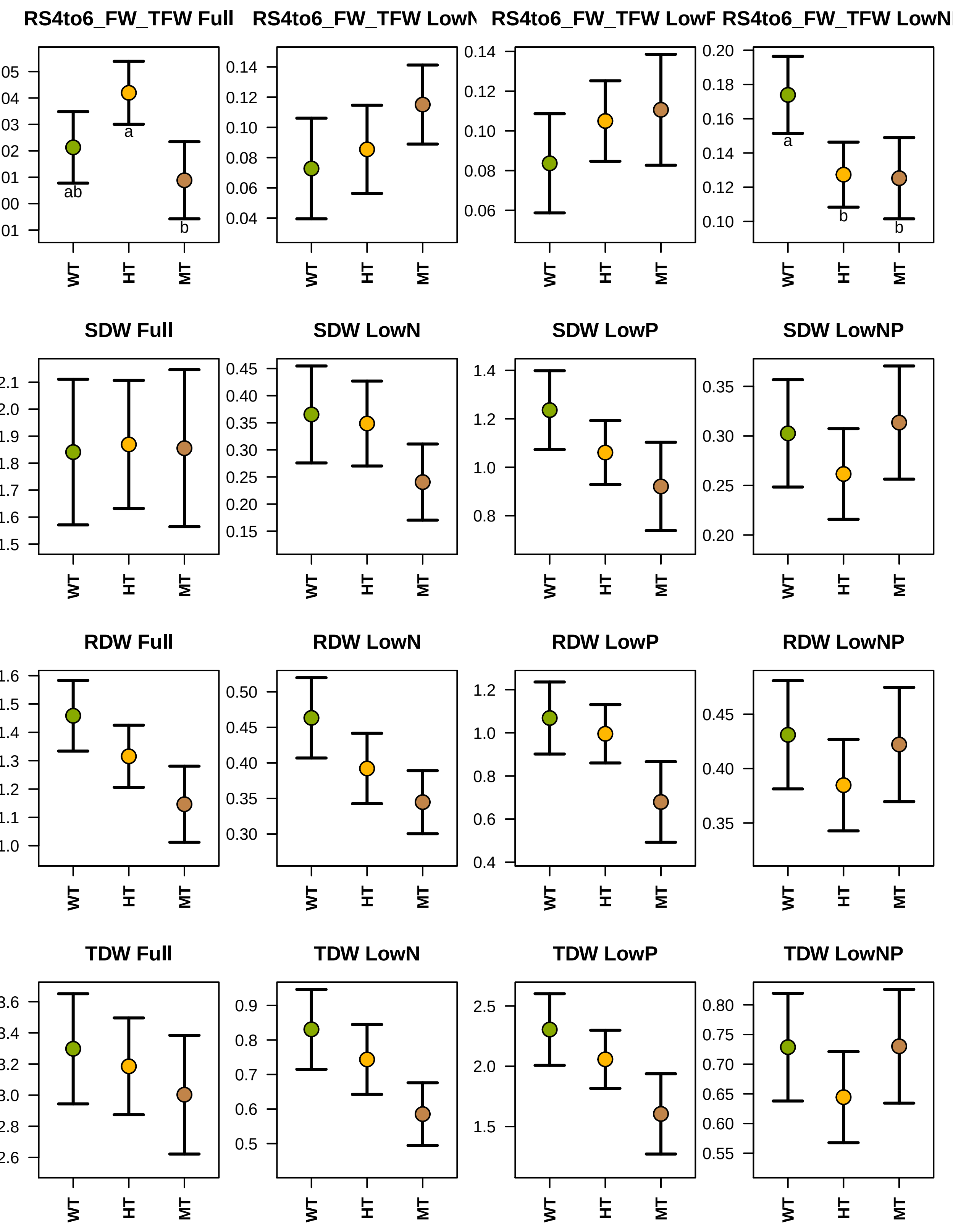

RS4to6_FW_TFW Full
RS4to6_FW_TFW LowN
RS4to6_FW_TFW LowP
RS4to6_FW_TFW LowNP
SDW Full
SDW LowN
SDW LowP
SDW LowNP
RDW Full
RDW LowN
RDW LowP
RDW LowNP
TDW Full
TDW LowN
TDW LowP
TDW LowNP
WT
HT
MT
ab
a
b

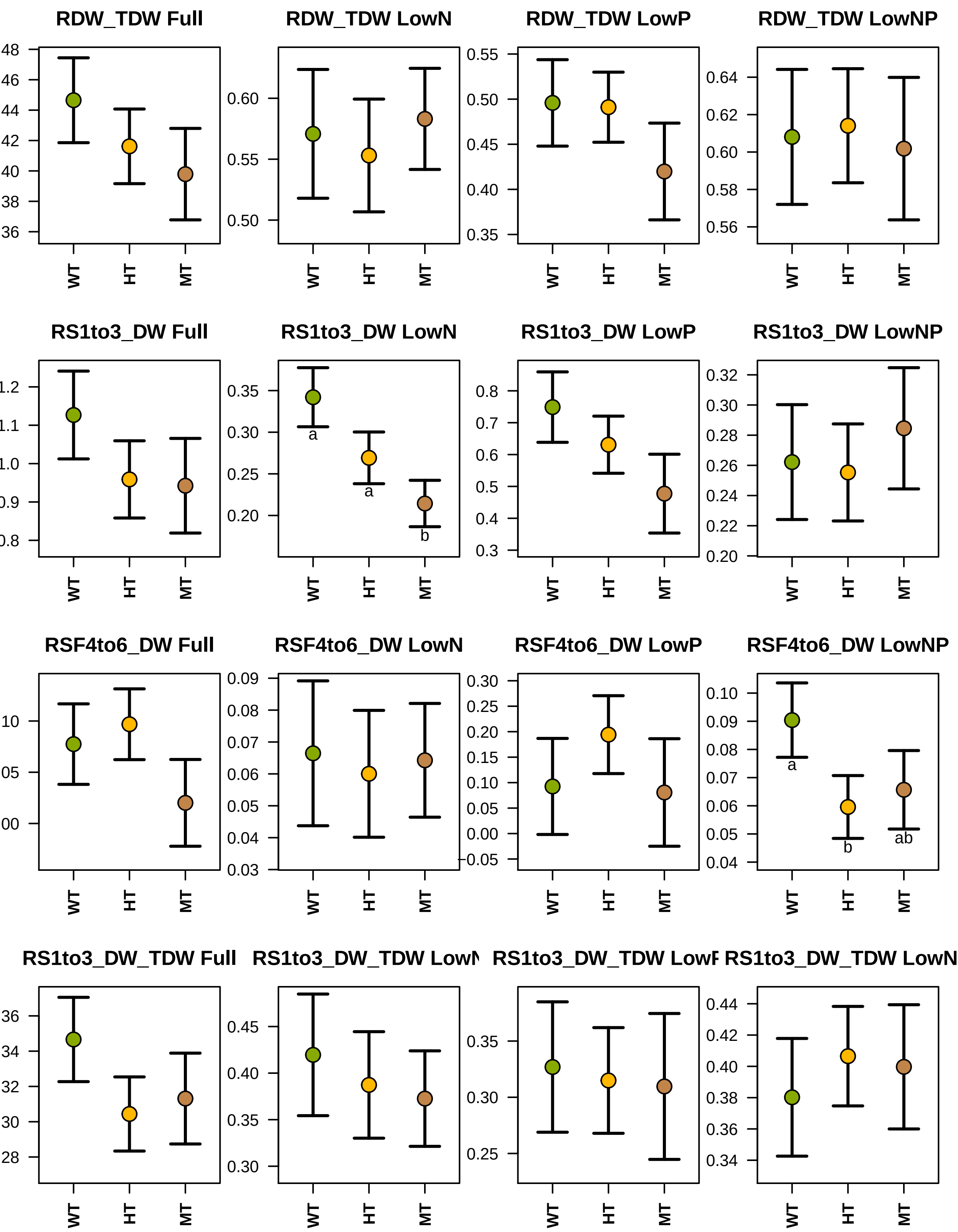
RDW_TDW Full
RDW_TDW LowN
RDW_TDW LowP
RDW_TDW LowNP
RS1to3_DW Full
RS1to3_DW LowN
RS1to3_DW LowP
RS1to3_DW LowNP
RSF4to6_DW Full
RSF4to6_DW LowN
RSF4to6_DW LowP
RSF4to6_DW LowNP
RS1to3_DW_TDW Full
WT
HT
MT
a
a
b
a
b
ab

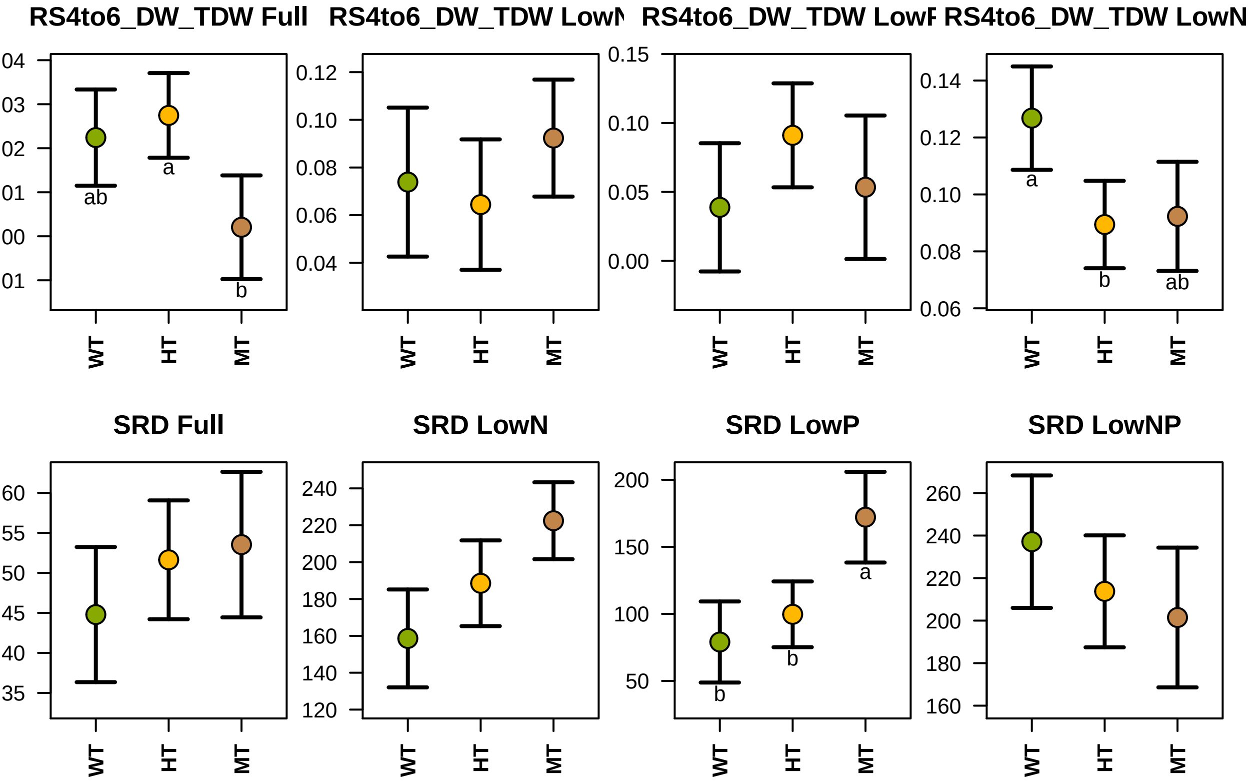
RS4to6_DW_TDW Full
RS4to6_DW_TDW LowN
RS4to6_DW_TDW LowP
RS4to6_DW_TDW LowN
ab
a
b
a
b
ab
b
b
a
0.12
0.10
0.08
0.06
0.04
0.15
0.10
0.05
0.00
0.14
0.12
0.10
0.08
0.06
WT
HT
MT
SRD Full
SRD LowN
SRD LowP
SRD LowNP
60
55
50
45
40
35
240
220
200
180
160
140
120
200
150
100
50
260
240
220
200
180
160
