## Supplemental Figure 3 for "The *pho1;2a’-m1.1* allele of *Phosphate1* conditions mis-regulation of the phosphorus starvation response in maize (*Zea mays* ssp. *mays* L.)"

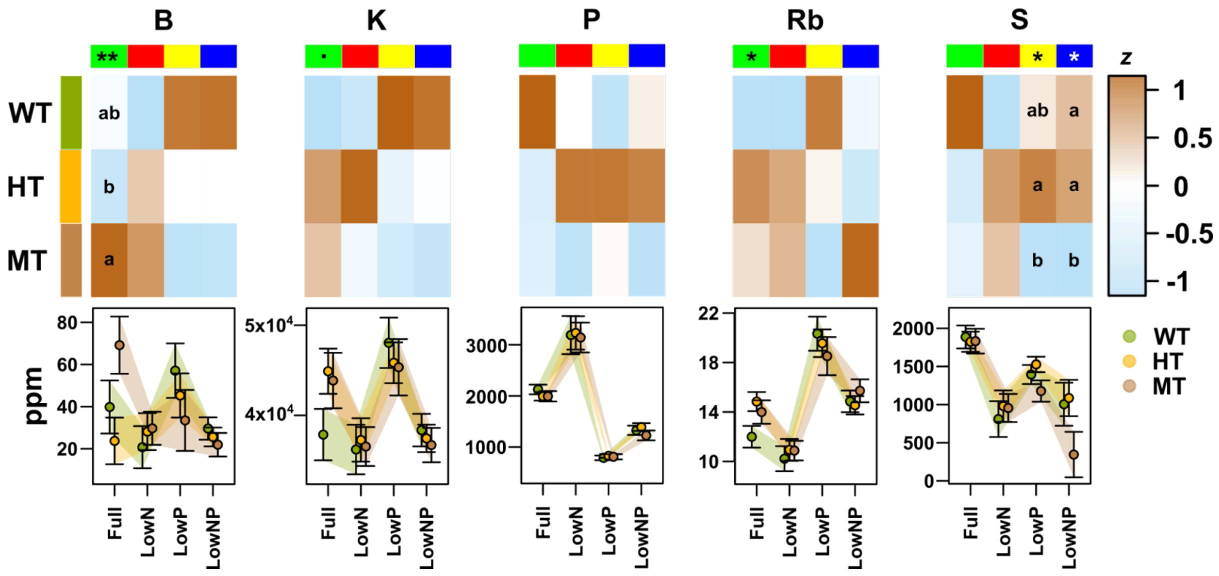


**Supplemental Figure 3. Leaf ionome nutrient responses are modulated in *pho1;2a´m1.1* mutants.** Total concentration of selected elements in the leaf displayed as heatmaps of standardized values (z value, standardized across genotypes for each element/treatment combination) and as plots (ppm). The genotype term was significant in the element/treatment combinations shown as *** $p < 0.001$, ** $p < 0.01$, * $p < 0.05$, . $p < 0.1$ (value before adjustment for multiple testing). Significant pairwise differences ($p < 0.05$) recovered from Tukey analysis are indicated with lowercase letters. Plots show estimated coefficients with bars extending +/- 1 standard error (SE).
