## Supplemental Figure 5 for "The *pho1;2a’-m1.1* allele of *Phosphate1* conditions mis-regulation of the phosphorus starvation response in maize (*Zea mays* ssp. *mays* L.)"

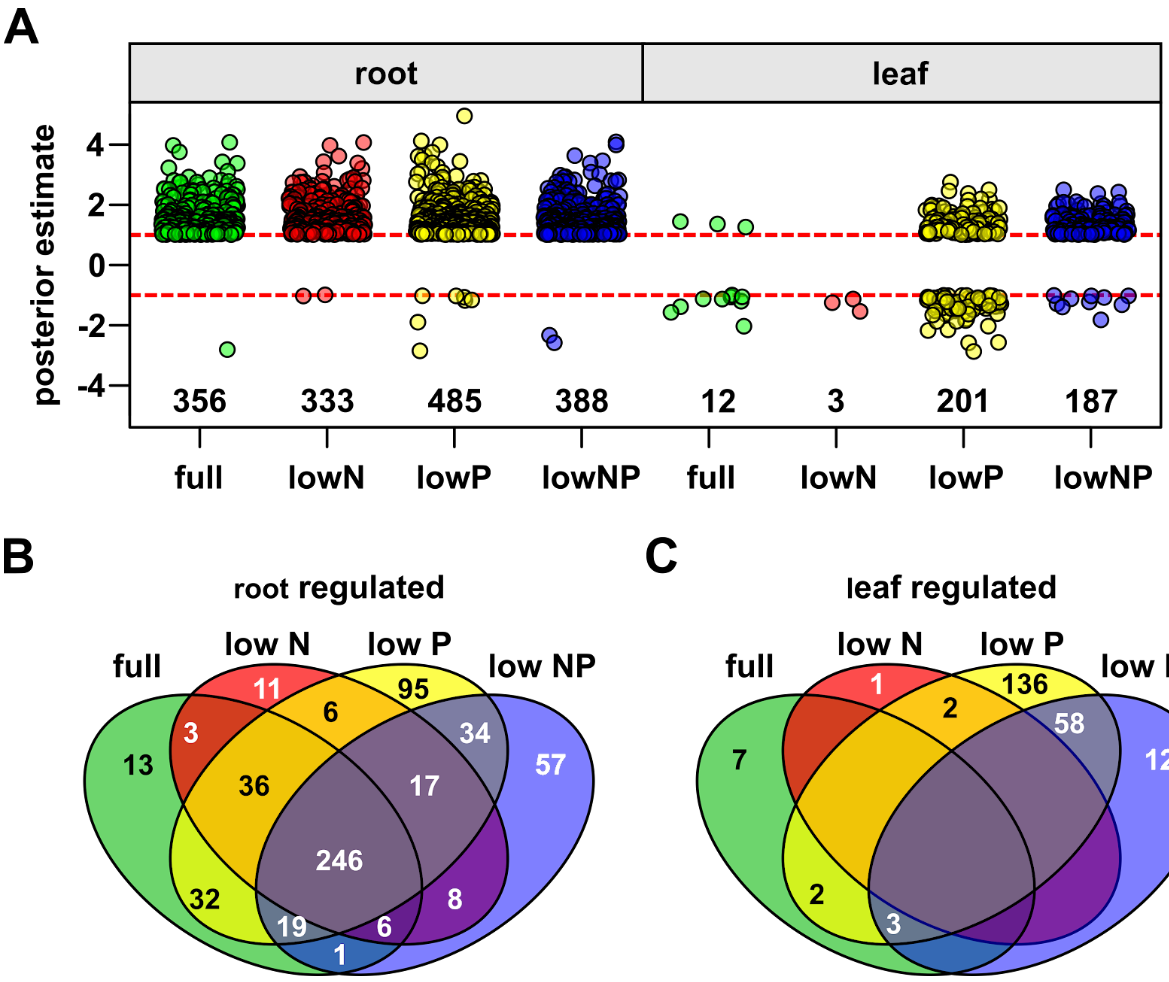


**Supplemental Figure 5. Plants heterozygous for the *pho1;2a´m1.1* mutation exhibit transcriptional mis-regulation.** A) Effect size estimates (log2 fold change; heterozygous - wild-type) for genes differentially expressed in plants heterozygous for *pho1;2a* compared with segregating wild-type siblings. Only significant genes are shown in any given tissue-treatment combination. Numbers at the base of the plot show the count of differentially expressed genes. A gene with an effect size >5 in the roots is not shown. B) Venn diagrams showing the overlap (count) of genes differentially expressed in the root (heterozygous *pho1;2a* compared with segregating wild-type siblings) among the four nutrient treatments. C) as B, for genes differentially expressed in the leaf.
